# Extensive diversity of RNA viruses in ticks revealed by metagenomics in northeastern China

**DOI:** 10.1101/2022.04.27.489762

**Authors:** Ziyan Liu, Liang Li, Wenbo Xu, Yongxu Yuan, Xiaojie Liang, Li Zhang, Zhengkai Wei, Liyan Sui, Yinghua Zhao, Yanyan Cui, Qing Yin, Dajun Li, Qianxue Li, Feng Wei, Zhijun Hou, Quan Liu, Zedong Wang

## Abstract

Recently, several emerging tick-borne viruses have been identified to be associated with human diseases in northeastern China. Here, we used metagenomics to investigate the virome diversity in *Haemaphysalis japonica, H. conicinna, Dermacentor silvarum*, and *Ixodes persulcatus* ticks in northeastern China. A total of 22 RNA viruses were identified and belonged to more eight viral families, including four each in *Nairoviridae* and *Phenuiviridae*, three each in *Flaviviridae, Rhabdoviridae*, and *Solemoviridae*, two in *Chuviridae*, and one each in *Partitiviridae, Tombusviridae*, and unclassified. Of them, eight viruses were novel species, belonging to *Nairoviridae* (Ji’an nairovirus and Yichun nairovirus), *Phenuiviridae* (Mudanjiang phlebovirus), *Rhabdoviridae* (Tahe rhabdovirus 1-3), *Chuviridae* (Yichun mivirus), and *Tombusviridae* (Yichun tombus-like virus), and five members were established human pathogens, including Alongshan virus, tick-borne encephalitis virus, Songling virus, Beiji nairovirus, and Nuomin virus. *I. persulcatus* ticks had significant higher viral species than those in *H. japonica, H. concinna*, and *D. silvarum* ticks. Significant differences in tick viromes were observed among Daxingan, Xiaoxingan and Changbai mountains. These findings showed an extensive diversity of RNA viruses in ticks in northeastern China, revealed potential public health threats from the emerging tick-borne viruses. Further studies are needed to explain the natural circulation and pathogenicity of these viruses.

## Introduction

Ticks are obligate haematophagous ectoparasites of wild and domestic animals as well as humans, and there are approximately 900 tick species worldwide, of which many can transmit pathogenic agents, including viruses, bacteria, and protozoa (1). Several tick-borne viruses (TBVs) are associated with serious diseases in humans and animals, such as tick-borne encephalitis virus (TBEV), Crimean-Congo hemorrhagic fever virus (CCHFV), and Nairobi sheep disease virus (NSDV) (2). In recent decades, the incidence and geographical distribution of tick-borne viruses have an increasing tendency, highlighting the public health importance of these arboviruses (3). Due to the application next generation sequencing (NGS) in recent years, many novel viruses have been identified in different tick species of different regions worldwide (4-7). To our knowledge, the known identified TBVs include hundreds of viral members of at least 12 genera in 9 families of two orders as well as other unassigned members (8).

In China, tick-borne encephalitis virus (TBEV) in the family *Flaviviridae* and Crimean-Congo hemorrhagic fever virus (CCHFV) in the family *Nairoviridae* are the two causative agents of viral encephalitis and hemorrhagic fever in northeastern and northwestern regions, respectively, that are transmitted by different tick species (9-11). Emerging TBVs, such as severe fever with thrombocytopenia virus (SFTSV), Jingmen tick virus (JMTV), Alongshan virus (ALSV), Songling virus (SGLV), Beiji nairovirus (BJNV), Tacheng tick virus 1 and 2, have been reported to be associated with human diseases (12-18). Nairobi sheep disease virus (NSDV), another member in the family *Nairoviridae* that can cause an acute hemorrhagic gastroenteritis in sheep and goats, has also been found in ticks in China (19, 20). Therefore, it is necessary to conduct routine surveillance of tick-borne viruses. The high-throughput sequencing technology has been widely used to investigate viromes in various species and to diagnose unknown infectious diseases. The tick viromes have been analyzed in several provinces, including Heilongjiang, Liaoning, Hebei, Henan, and Yunnan in China (21-25), revealing a large number of novel RNA viruses of vertebrate and invertebrate hosts. These studies also suggest that the viromes are significantly affected by the tick species and geographical location.

The northeastern region has the richest forest resources in China, mainly concentrated in the Daxingan mountain (DXAM), Xiaoxingan mountain (XXAM) and Changbai mountain (CBM), leading to abundant tick populations. In addition to TBEV, several emerging tick-borne viruses that may infect humans have been discovered in northeastern China, such as ALSV (14), SGLV (26), BJNV (16), and JMTV (13). Here, using metagenomic analysis, we found extensive diversity of RNA viruses in ticks in northeastern China, revealing potential public threats from the emerging TBVs. Further studies are needed to explain the natural circulation and pathogenicity of these viruses.

## 2. Materials and methods

### 2.1 Sample collection

The questing ticks were collected by the flagging method, and blood-sucking ticks were collected from cattle in Shulan. These ticks were identified to species based on the morphological criteria combined with polymerase chain reaction (PCR) of 16S ribosomal RNA gene as described elsewhere (27, 28). Every 10 ticks were pooled into a 1.5 mL Eppendorf tube according to the collection sites and species and stored at -80°C until further study.

### 2.2 RNA library construction and sequencing

After washing with 75% ethanol and RNA/DNA-free water, pooled ticks in tubes were added with 800 μL Dulbeccos modified Eagles minimum essential medium (DMEM) and two stainless steel beads (3 mm diameter), and crushed using the Tissuelyser (Jingxin, Shanghai, China) at 70 Hz for 2 min. The lysates were centrifuged at 12000 rpm for 10 min at 4°C, and the supernatant was collected for viral RNA extraction with the TIANamp Virus RNA kit (TIANGEN, Beijing, China). The viral RNA was further pooled for library construction according to the collection sites and species (Supplementary Table S1).

The pooled RNA was digested with micrococcal nuclease (NEB, USA) in 37°C for 2 h, followed by metagenomic sequencing at Tanpu Biological Technology Co., LTD (Shanghai, China). Briefly, the RNA from each pool was used for library preparation with the NEBNext® UltraTM RNA Library Prep Kit for Illumina® (NEB, USA) according to the manufacturer’s instructions. After adapter ligation, ten cycles of PCR amplification were performed for sequencing target enrichment. The libraries were pooled at equal molar ratio, denatured and diluted to optimal concentration, and sequenced with an Illumina NovaSeq 6000 System.

### 2.3 Transcriptome analysis

Transcriptome analysis was conducted as described elsewhere (29). Briefly, after trimming and removing low quality reads, the paired-raw reads were purified by removing ribosomal RNA, host contamination, and bacteria sequences using BBMap program (https://github.com/BioInfoTools/bbmap), and assembled into contigs with SPAdes v3.14.1 (https://github.com/ablab/spades) and SOAPdenovo v2.04 (https://github.com/aquaskyline/SOAPdenovo-Trans) (30, 31). After being compared with the non-redundant nucleotide (nt) and protein (nr) database downloaded from GenBank using BLAST+ v2.10.0, the assembled contigs were filtered to remove the host and bacterial sequences. The relative abundance of the identified viruses was determined by mapping the reads back to the assembled contigs using Bowtie2 v2.3.3.1.

### 2.4 Viral genome confirmation and annotation

The assembled contigs were compared with NCBI nucleotide and viral refseq database using BLAST (V2.10.0+), and used as a reference for designing specific primers to confirm and analyze the sequences of terminal ends, using the nested reverse transcription-polymerase chain reaction (RT-PCR) and the rapid amplification of cDNA ends (RACE) performed as described elsewhere(16, 26). Detailed information on the primers for the detection or whole genome amplification are shown in Supplementary Table S2. Potential open reading frames (ORFs) in the viral sequences were predicted using ORFfinder (https://www.ncbi.nlm.nih.gov/orffinder/).

### 2.5 Virus classification

All the viruses identified in this study were classified according to the latest International Committee on Taxonomy of Viruses (ICTV) report of virus taxonomy (https://talk.ictvonline.org/ictvreports/ictv_online_report/). A novel viral species should be satisfied with one of the following conditions as described before (22), namely, (i) <80 per cent nt identity across the complete genome; or (ii) <90 per cent aa identity of the RNA-dependent RNA polymerase (RdRp) domain with the known viruses. All novel viruses were named as the collection sites that the virus was first identified, followed by common viral names according to their taxonomy. All the viral strains would be marked with ‘Northeatern (NE)’ to distinguish them from the virus strains identified in other studies.

### 2.6 Phylogenetic analyses

To confirm the phylogenetic relationships of the viruses discovered in this study, representative reference sequences were retrieved from GenBank database (Supplementary Table S3), and the nucleotide (nt) and amino acid (aa) sequences were aligned using ClustalW available within MEGA 7.0. Phylogenetic analyses were conducted with the aligned sequences using the maximum-likelihood method in MEGA version 7.0 with the best-fit substitution model for each alignment (32). A bootstrapping analysis of 1000 replicates were conducted in the analysis, and the bootstrap values more than 70 were shown in the trees.

## 3. Results

### 3.1 Tick collection and identification

From April 2020 to July 2021, a total of 2,031 ticks, including 204 *Haemaphysalis japonica*, 393 *H. conicinna*, 386 *Dermacentor silvarum*, and 1048 *Ixodes persulcatus* ticks, were collected from NE China (Fig.1, Supplementary Table S1). The collection sites were distributed in Ji’an (n=100), Dunhua (n=382), and Shulan (n=300) in Jilin Province, Fangzheng (n=300), Mudanjiang (n=156), Yichun (n=253), and Tahe (n=316) in Heilongjiang Province, and Songling (n=224) in Inner Mongolia Autonomous Region (Fig.1, Supplementary Table S1). Of these sampling sites, five were located in Changbai Mountain (CBM), two in Daxingan Mountain (DXAM), and one in Xiaoxingan Mountain (XXAM) (Fig.1, Supplementary Table S1).

**Figure 1.**
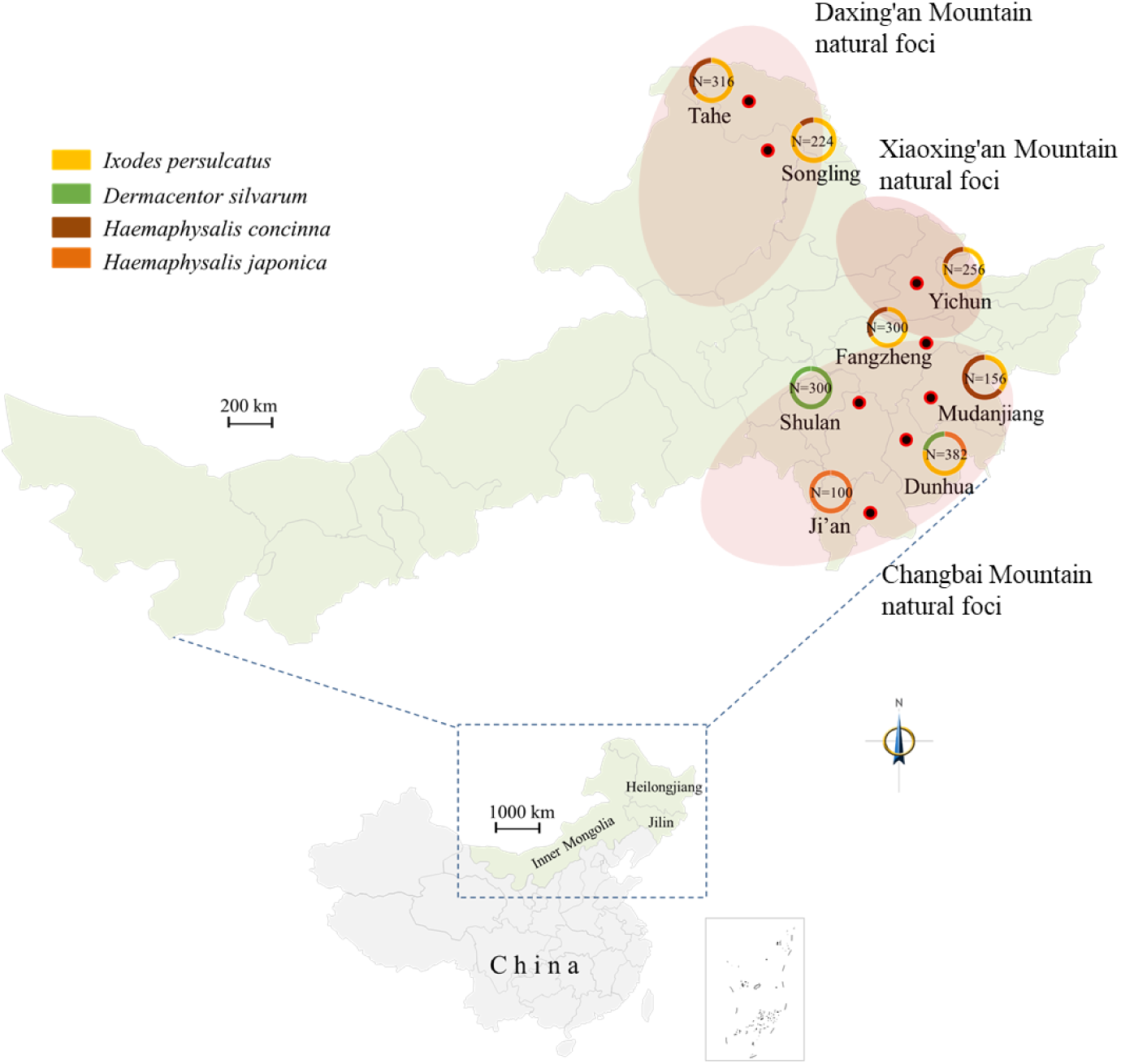
Sampling sites of ticks in northeastern China. The number of ticks collected at each site was marked, and species was shown by different colors. Yellow, *Ixodes persulcatus*; Green, *Dermacentor silvarum*; Brown, *Haemaphysalis conicinna*; Orange, *Haemaphysalis japonica*.

### 3.2 Identified RNA viruses

A total of 24 RNA libraries were constructed and sequenced, resulting to 245.8 GB clean data and ∼0.5 billion non-rRNA reads (Supplementary Table S1). Totally, 2,059 viral contigs were obtained by de novo assembly from ∼0.93 million viral reads that accounted for 0.2% of the total non-rRNA reads. Within each library, the viral reads ranged from 0.02% (library SL3) to 0.53% (library JA1) of the total non-rRNA reads. After being aligned by Blast and filtered by virus-host database, the viral contigs were finally annotated to 22 viruses, belonging to the 8 viral families, including *Flaviviridae, Rhabdoviridae, Nairoviridae, Phenuiviridae, Chuviridae, Partitiviridae, Tombusviridae, Solemoviridae*, and one unclassified viral species (Table 1).

**Table 1.** Viruses identified in the present study.

| Classification | Virus (abbreviation) | Genome (bp) | Closet relative (% nt identity) | No.<sup>a</sup> |
| --- | --- | --- | --- | --- |
| **Flaviviridae** |  |  |  |  |
| Jingmenvirus | Alongshan virus (ALSV) | 3,066/2,813/2,809/2,721 | Alongshan virus H3 (96.9, 98.3, 98.0, 98.6) | 1 |
| Flavivirus | Tick-borne encephalitis virus (TBEV) | 11,048 | Tick-borne encephalitis virus HLB-T74 (97.8) | 3 |
| Pestivirus-like | Bole tick virus 4 (BLTV-4) | 16,281 | Bole tick virus 4 Iasi20 (85.3) | 2 |
| **Nairoviridae** |  |  |  |  |
| Orthonairovirus | Songling virus (SGLV) | 12,030/4,525/2,132 | Songling virus YC585 (95.9, 98.2, 98.5) | 2 |
| Orthonairovirus | Ji'an nairovirus (JANV) | 11,993/4,404/1,802 | Songling virus YC585 (73.7, 73.1, 71.9) | 4 |
| Norwavirus-like | Beiji nairovirus (BJNV) | 14,899/3,767 | Beiji nairovirus H160 (98.97, 98.6) | 7 |
| Norwavirus-like | Yichun nairovirus (YCNV) | 14,813/3,326 | Beiji nairovirus H1063 (83.0, 80.9) | 3 |
| ***Phenuiviridae*** |  |  |  |  |
| *Phlebovirus* | Mukawa virus (MKWV) | 6,422/3,309/1,887 | Mukawa virus MKW73 (92.2, 92.2, 93.5) | 6 |
| *Phlebovirus* | Mudanjiang phlebovirus (MJPV) | 6,428/3,330/1,882 | Kuriyama virus CZCT80Q (77.5, 70.9, 72.5) | 2 |
| *Ixovirus* | Sara tick phlebovirus (STPV) | 6,706/2,535 | Sara tick phlebovirus Rus/Ix_persulcatus/Karelia/4/2018 (98.1, 98.8) | 6 |
| *Ixovirus* | Onega tick phlebovirus (OTPV) | 6,689/1,935 | Onega tick phlebovirus Rus/Ix_persulcatus/Karelia/3/2018 (98.3, 98.3) | 6 |
| ***Rhabdoviridae*** |  |  |  |  |
| Alphanemrha virus-like | Tahe rhabdovirus 1 (THRV1) | 11,343 | Manly virus (67.0) | 8 |
| Alphanemrha virus-like | Tahe rhabdovirus 2 (THRV2) | 11,486 | Norway mononegavirus 1 NOR/H3/Skanevik/2014 (67.0) | 2 |
| Alphanemrha virus-like | Tahe rhabdovirus 3 (THRV3) | 10,375 | Norway mononegavirus 1 NOR/H3/Skanevik/2014 (65.2) | 1 |
| ***Chuviridae*** |  |  |  |  |
| *Mivirus* | Nuomin virus (NUMV) | 10,900 | Nuomin virus Rus/Ix_persulcatus/Karelia/5/2018 (97.5) | 10 |
| *Nigecruvirus* | Yichun mivirus (YCMV) | 11,493 | Blacklegged tick chuvirus-2 RTS126 (72.6) | 2 |
| ***Tombusviridae*** |  |  |  |  |
| Tombusvirus-like | Yichun tombus-like virus (YTLV) | 4,369 | Upmeje virus OTU9. IU20 (74.39) | 4 |
| ***Solemoviridae*** |  |  |  |  |
| Sobemo-like | Jilin luteo-like virus 2 (JLLV2) | 2,731 | Jilin luteo-like virus 2 JL-QG-2 (99.1) | 6 |
| Sobemo-like | Xinjiang tick associated virus 1 (XTAV1) | 2,618 | Xinjiang tick associated virus 1 14-YG (96.1) | 4 |
| Sobemo-like | *Ixodes scapularis* associated virus 1 (ISAV1) | 2,658 | *Ixodes scapularis* associated virus 1 ISE6 (82.5) | 6 |
| ***Partitiviridae*** |  |  |  |  |
| Deltapartitivirus-like | Jilin partiti-like virus 1 (JPLV1) | 1,717 | Jilin partiti-like virus 1 JL/QG-4 (99.7) | 7 |
| **Unclassified** | *Ixodes scapularis* associated virus 3 (ISAV3) | 1,523 | *Ixodes scapularis* associated virus 3 ISE6 (84.6) | 5 |

<sup>a</sup> Number of RNA library with the indicated virus that was confirmed by RT-PCR and Sanger sequencing.

### 3.2 Viral genomic organization and taxonomy

For each of the 22 RNA viruses, the whole genomes of the representative viral strains were further verified by Sanger sequencing (Table 1 and Supplementary Fig. S1). Totally, 150 viral sequences from the 22 identified RNA viruses were verified and submitted to NCBI with the accession numbers showed in Supplementary Table S4. Of them, 13 viruses were found in China for the first time, and eight viruses were proposed as novel viral species, as they were highly divergent to any of previously identified viruses (nt identities < 80% or RdRp aa identities <90%), designated Tahe rhabdovirus 1-3 (THRV 1-3), Ji’an nairovirus (JANV), Yichun nairovirus (YCNV), Mudanjiang phlebovirus (MJPV), Yichun mivirus (YCMV), and Yichun tombus-like virus (YTLV), respectively (Table 1). Another eight viruses, including Alongshan virus (ALSV), Bole tick virus 4 (BLTV-4), Beiji nairovirus (BJNV), Jilin luteo-like virus 2 (JLLV2), Xinjiang tick associated virus 1 (XTAV1), *Ixodes scapularis* associated virus 1 (ISAV1), Jilin partiti-like virus 1 (JPLV1), and *Ixodes scapularis* associated virus 3 (ISAV3), showed the close relationships (nt identities >80 per cent and RdRp aa identities>90 per cent) with previously described tick-associated viruses that have not been approved by ICTV yet (Table 1). The remaining six viruses were all classified into known species, as they showed close relationships and identical genome organizations with ICTV-approved viruses, including tick-borne encephalitis virus (TBEV), Songling virus (SGLV), Mukawa virus (MKWV), Sara tick phlebovirus (STPV), Onega tick phlebovirus (OTPV), and Nuomin virus (NUMV) (Table 1).

### 3.3 Virome composition and abundance

Across the 24 libraries, 22 possessed 1-11 viral species, with the exception of the libraries FZ1 and YC1 that had no virus detectable (Fig. 2A). The ten libraries of *I. persulcatus* ticks had 4-11 viral species, with 11 species in the library YC4 that included BJNV, YCNV, MKWV, STPV, OTPV, NUMV, YCMV, JPLV1, JLLV2, ISAV1, and ISAV3. In contrast, *H. japonica, H. concinna*, and *D. silvarum* libraries only possessed 0-3 viral species. Of these 22 viral species, 17 species were identified in *I. persulcatus*, while only two, three, and four species were detected in *H. japonica, H. concinna*, and *D. silvarum* ticks, respectively (Fig. 2B). Notably, most viruses were only identified in one tick species; however, Tahe rhabdovirus 1 was confirmed in the *H. japonica, H. concinna*, and *D. silvarum* ticks, Ji’an nairovirus was detected in the *H. japonica* and *H. concinna* ticks, and Mukawa virus was identified in *D. silvarum* and *I. persulcatus* ticks. It should also be noted that both ALSV and THRV3 were detected in only one *I. persulcatus* library. Interestingly, some viral species, such as THRV2-3, SOLV, TBEV, and ALSV, were only detected in ticks of DXAM, while other virus, including JANV, MDPV, YCNV, YCMV, BLTV4, YTLV, and XTAV1, were detected in ticks from XXAM and CBM. There were also some viral species detectable in the libraries of three regions, such as THRV1, BJNV, MKWV, NUMV, JPLV1, JLLV2, and ISAV1, showing the wide distribution of these viruses (Fig. 2B). Of them, the libraries of *I. persulcatus* ticks had relatively higher viral reads than those in *H. japonica, H. concinna*, and *D. silvarum* ticks, and the libraries of XXAM (Yichun) and DXAM (Tahe and Songling) had relatively higher viral reads than that in the libraries of CBM. Moreover, NUMV and BJNV had relatively higher viral reads than other viral species (Fig. 2B).

**Figure 2.**
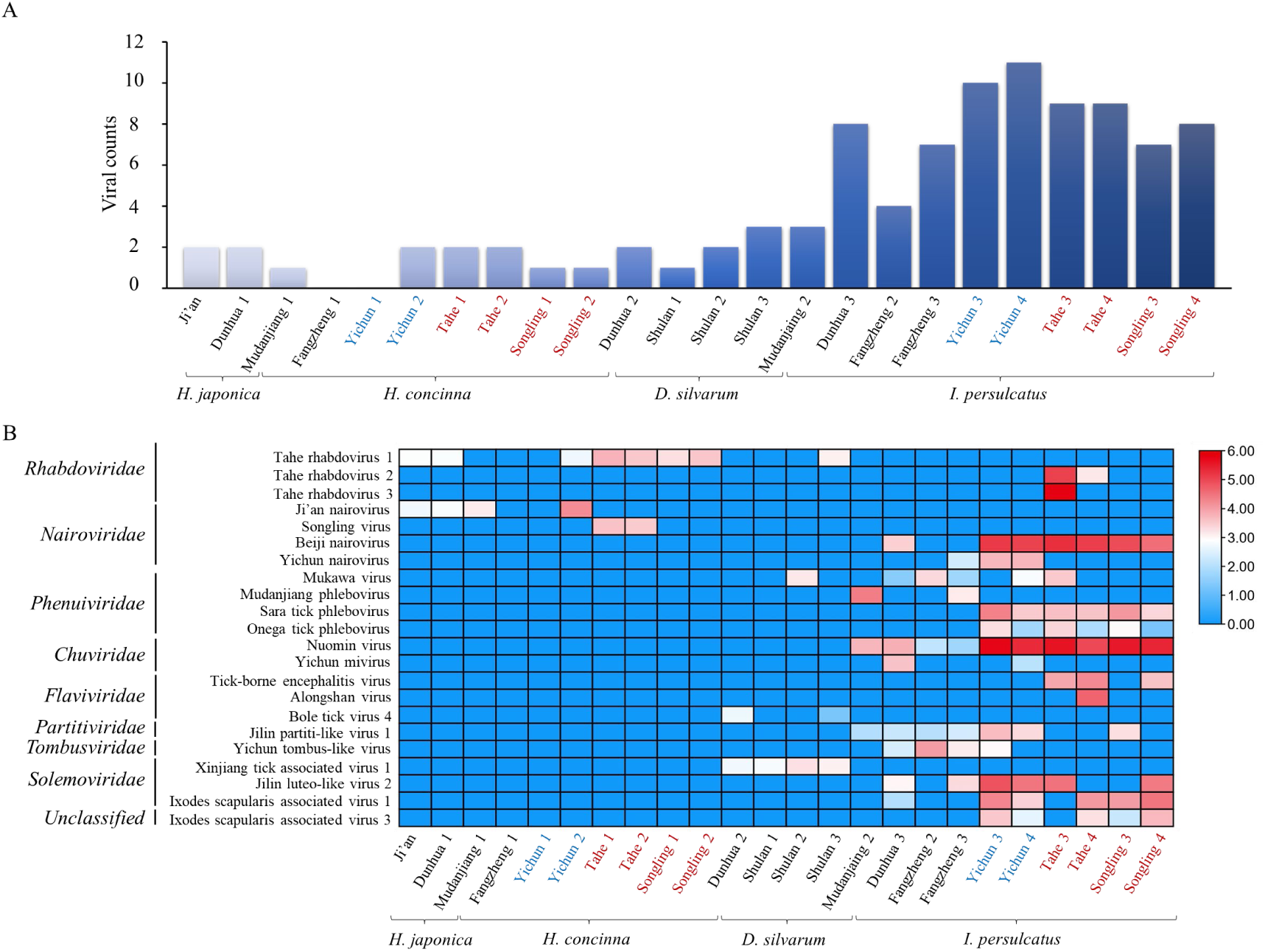
Viral presence and abundance across the libraries. (A) Viral number identified in each library. (B) Heatmap based on the normalized numbers of sequence reads for 8 viral families and one unclassified RNA virus in each library. Tick species and location information were listed at the bottom, and names of the viral families and species were indicated on the left. Log10 relative abundance of the viruses in each species in each location were indicated as a heat map ranging from low (blue) to high (red) based on the normalized average viral genome size and total sequencing reads in each library. The libraries from XXAM, DXAM, and CBM are marked with blue, red, and black, respectively.

#### 3.4.2.1 Flaviviridae

According to the latest ICTV report of virus taxonomy, *Flavivirus, Hepacivirus, Pegivirus*, and *Pestivirus* are the approved genera in family *Flaviviridae*. While in recent years, a flavivirus-like group, the Jingmenvirus group, including a series of genetically related viruses, such as Jingmen tick virus, ALSV, and Yanggou tick virus, have been discovered (Fig. 3A). In this study, ALSV was identified in the *I. persulcatus* library (TH4) from Tahe in DXAM, and clustered together with ALSV strain H3 isolated from tick-bitten patients in NE China (Fig. 3B) (14), but different from the isolates detected in *I. persulcatus* in Russia and *I. ricinus* in Finland (Fig. 3B), with nt identities of 97.0–98.5% (Extended data Table E1, 2) (33, 34).

**Figure 3.**
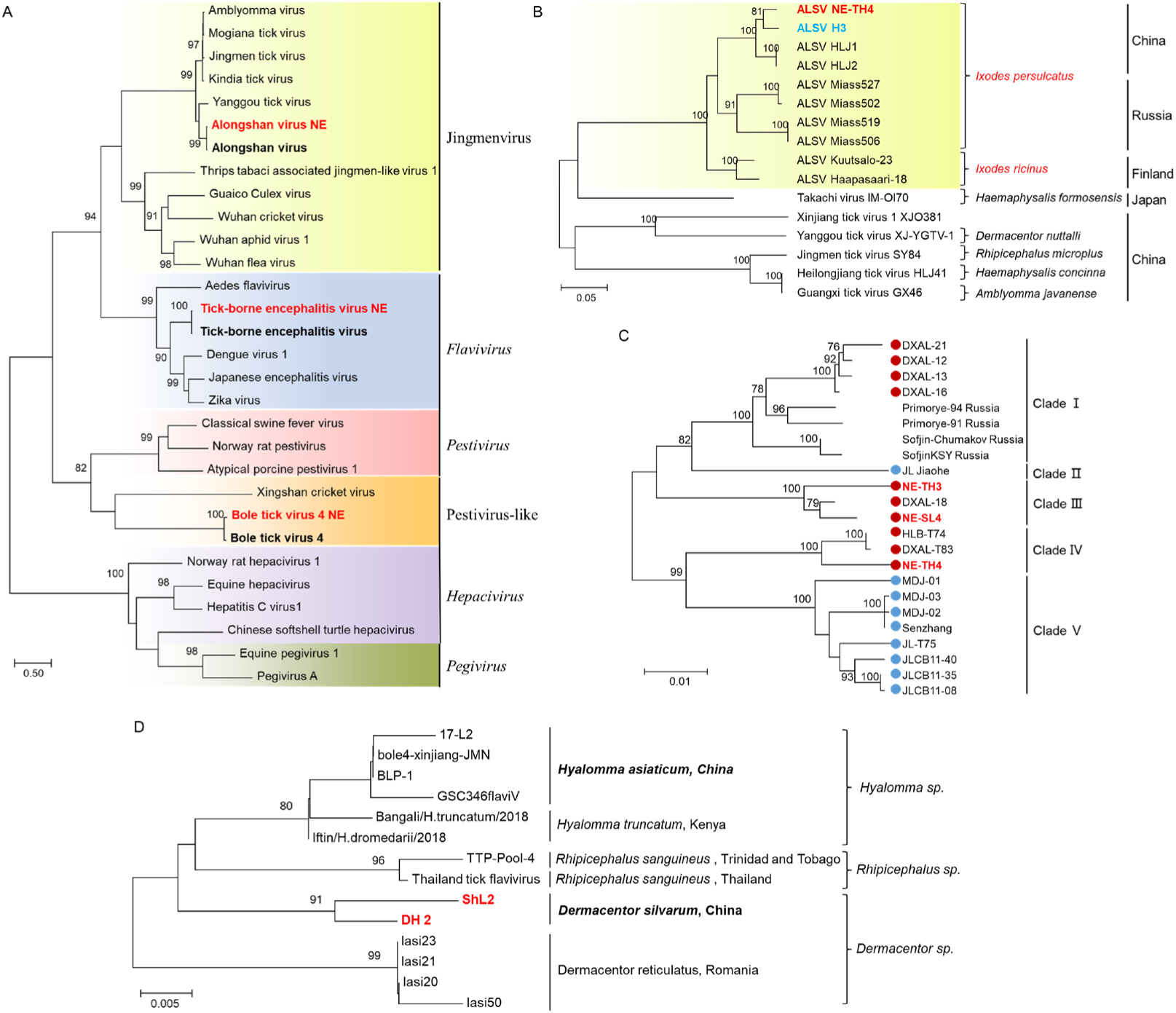
Phylogenetic analyses of flaviviruses. Phylogenetic trees were constructed based on the RdRp sequence of representative viruses in the family *Flaviviridae* (A), the segment 3 of ALSV and other representative viral strains in the Jingmenvirus group (B), the E protein of TBEV in NE China (C), and the RdRp of BLTV4 (D). All the viruses obtained in ticks here were highlighted in red. In panel A, closest referenced viruses were highlighted in bold font, and the virus genera or groups were marked with different colors of background. In panel B, ALSV isolated from humans were marked with blue. ALSV were highlighted with light green background. The host tick species of the viruses and the countries that the viruses discovered were also labeled. In panel C, TBEV isolated from CBM and XXAM were marked with blue-filled circles, while strains found in DXAM were marked with red-filled circles. In panel D, the strain names of BLTV4 were labeled. The host tick species and the countries that BLTV4 discovered were also marked. Abbreviations: ALSV, Alongshan virus; TKCV, Takachi virus; XJTV, Xinjiang tick virus; YGTV, Yanggou tick virus; HLJTV, Heilongjiang tick virus; JMTV, Jingmen tick virus; GXTV, Guangxi tick virus; TBEV, Tick-borne encephalitis virus; BLTV4, Bole tick virus 4. The accession numbers of the viral sequences used in the trees were shown in Supplementary Table S3, 4.

Three libraries (TH3, TH4, and SL4) of *I. persulcatus* ticks from Tahe and Songling in DXAM were identified TBEV-positive, which formed a different clade from the TBEV strains in XXAM and CBM, with nt identities of 93.5–99.9% (Fig. 3C, Extended data Table E3).

Bole tick virus 4 (BLTV4)-NE strains were phylogenetically grouped into the pestivirus-like group, with nt identities of 77.5–83% and RdRp aa identities of 94.6–96.9% to other BLTV4 isolates (Fig. 3A, Extended data Table E4). BLTV4 was first identified in *Hyalomma asiaticum* ticks in Xinjiang Uygur Autonomous Region, China (35), and also discovered in Trinidad and Tobago (36), Kenya (37), Romania (38), and Thailand (39). Phylogenetic analysis showed that BLTV4-NE was different from other strains identified in different tick species or regions (Fig. 3D). In this study, the virus was only detected in two libraries of the *D. silvarum* ticks from CBM.

#### 3.4.2.2 Nairoviridae

In the phylogenetic tree of the family *Nairoviridae*, SGLV-NE, together with SGLV, formed a separate clade from other viral members in the genus *Orthonariovirus*, including Ji’an nariovirus, Tacheng tick virus 1, and Tamdy virus (Fig. 4A). SGLV-NE was identified in two libraries TH1 and TH2 of the *H. concinna* ticks from Tahe in DXAM, and clustered together with SGLV strains isolated from tick-bitten patients (Fig. 4B), with nt identities of 92.4–99.2% (Extended data Table E5, 6) (26).

**Figure 4.**
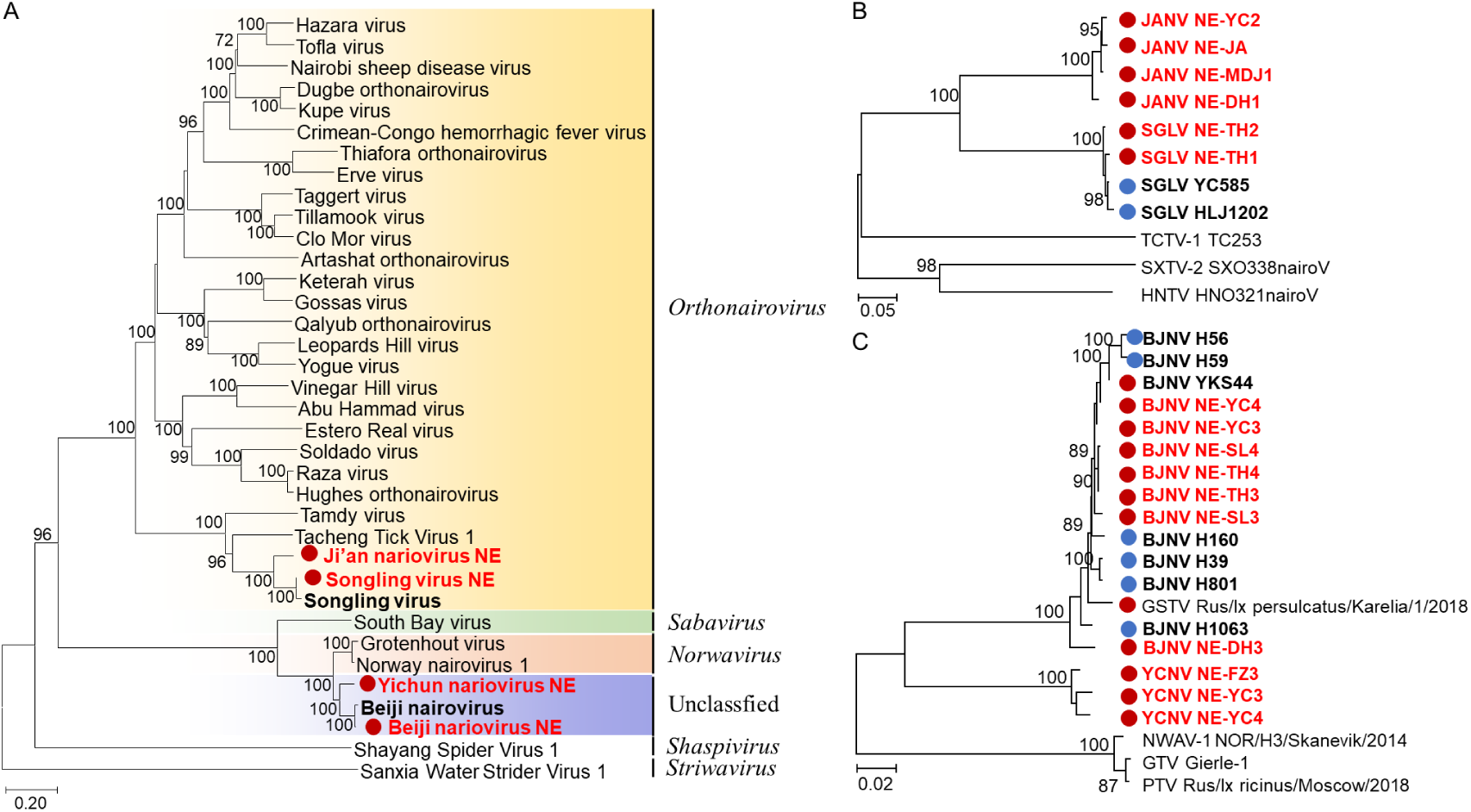
Phylogenetic analyses of nairoviruses. Phylogenetic trees were constructed based on the RdRp protein sequences of representative viruses in the family *Nairoviridae* (A), the S segment of JANV and SGLV (B), and the S segment of BJNV and YCNV (C). All the viruses obtained in ticks here were highlighted in red, and the closest referenced viruses were also highlighted in bold font. In panel B and C, viral strains discovered from ticks were marked with red-filled circles, while strains found in humans were marked with blue-filled circles. Abbreviations: JANV, Ji’an nariovirus; SGLV, Songling virus; TCTV1, Tacheng tick virus 1; SXTV2, Shanxi tick virus 2; HNTV, Henan tick virus; YCNV, Yichun nariovirus; BJNV, Beiji nariovirus; GKTV, Gakugsa tick virus; PTV, Pustyn virus; NWNV1, Norway nairovirus 1; GTV, Grotenhout virus. The accession numbers of the viral sequences used in the trees were shown in Supplementary Table S3, 4.

JANV, genetically related to SGLV with nt identities of 70.7-73.5%, was a novel identified nairovirus (Table S5, S6). Four libraries were detected JANV-positive, including DH1 and JA libraries of *H. japonica* ticks in CBM, and MDJ1 and YC2 of *H. concinna* ticks in XXAM (Fig. 4B).

BJNV and YCNV, belonging to an unclassified Norwavirus-like group, showed close relationships to Norway nairovirus 1 and Grotenhout virus (Fig. 4A), with nt identities of 75.4– 79.6% (Segment L and S, Extended data Table E7, 8). BJNV was identified in seven *I. persulcatus* tick libraries in all the three regions, suggesting the wide distribution of the virus in NE China. All the BJNV NE strains were clustered together with BJNV strains identified in tick-bitten patients and *I. persulcatus* tick in NE China and GSTV detected in Russia (Fig. 4C), with nt identities of 96.2–100% (Segment L and S, Extended data Table E7, 8).

YCNV was only detected in three libraries (FZ3, YC3, and YC4) of *I. persulcatus* ticks from Fangzheng and Yichun in CBM and XXAM. Although YCNV showed nt identities of 82.3–83.9 (>80) (Segment L and S, Extended data Table E7, 8) with BJNV, the virus formed a separate clade, with aa (RdRp) identities of 87.7–88.5% (Fig. 4C, Extended data Table E7, 8), indicating that YCNV was a novel viral species that may be different from BJNV.

#### 3.4.2.3 Phenuiviridae

MKWV and MJPV, together with STPV and OTPV identified here, fell within the genera *Phlebovirus* and *Ixovirus* of the family *Phenuiviridae*, respectively (Fig. 5A). MKWV NE strains were clustered together with the strain MKW73 identified in *I. persulcatus* ticks in Japan, with nt identities of 92.4–100% for L segment, 90.7–100% for M segment, and 93.5–99.9% for S segment (Fig. 5B, Extended data Table E9, 10). Of the six MKWV NE strains, five were detected in the *I. persulcatus* ticks from Shuangzi, Dunhua, Yichun, and Tahe, while one was identified in the *D. silvarum* ticks collected from cattle in Shulan.

**Figure 5.**
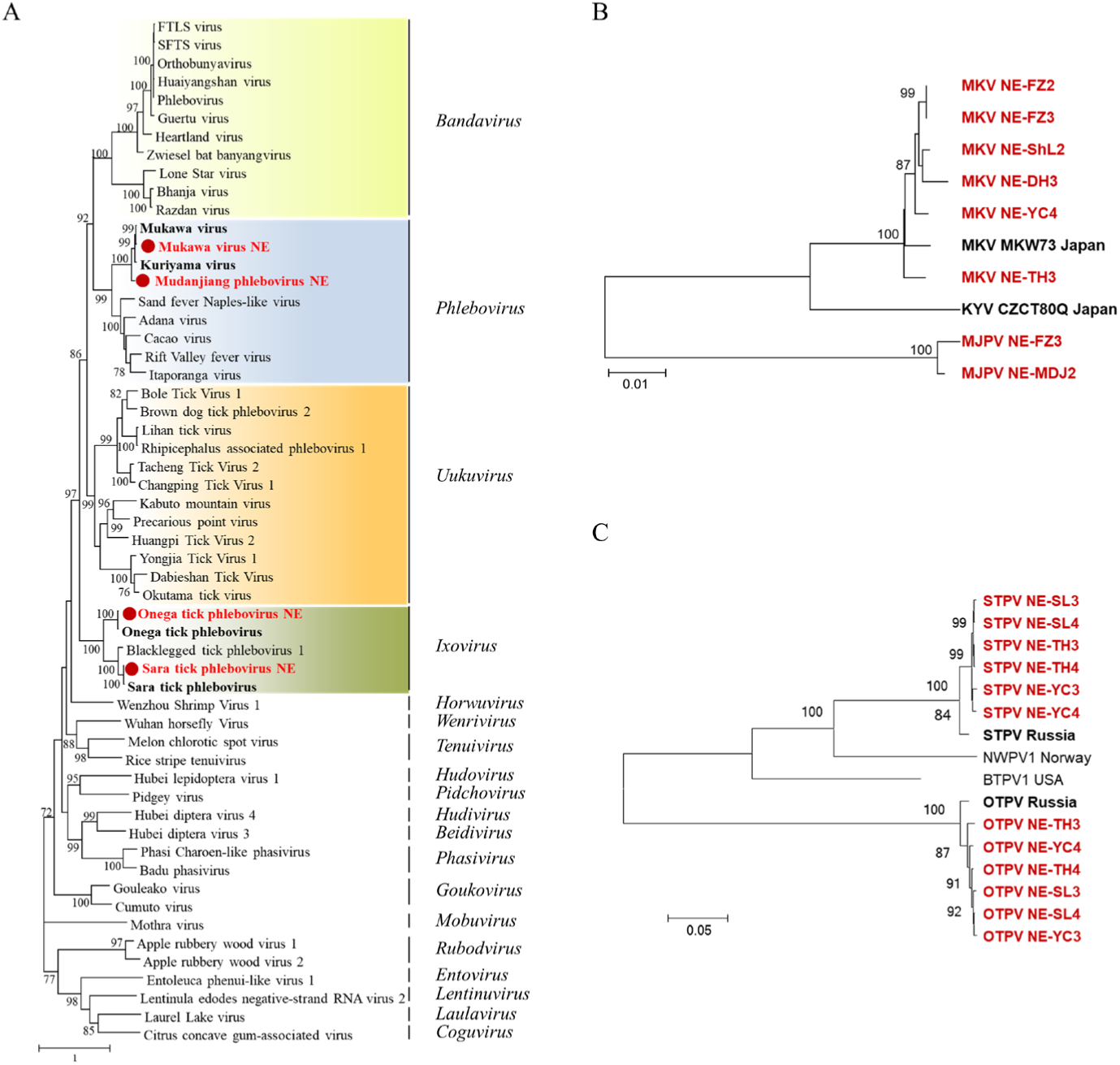
Phylogenetic analyses of phleboviruses. Phylogenetic trees constructed based on the RdRp sequences of representative viruses in the family *Phenuiviridae* (A), the S segment of MKWV and MJPV (B), and the S segment of STPV and OTPV (C). All the viruses obtained in ticks here were highlighted in red, and the closest referenced viruses were highlighted in bold font. Abbreviations: MKWV, Mukawa virus; KYV, Kuriyama virus; MJPV, Mudanjiang phlebovirus; BTPV1, Blacklegged tick phlebovirus-1; NWPV1, Norway phlebovirus 1; STPV, Sara tick phlebovirus; OTPV, Onega tick phlebovirus. The accession numbers of the viral sequences used in the trees are shown in Supplementary Table S3 and S4.

MJPV was only identified in two libraries in the *I. persulcatus* ticks from Fangzheng and Mudanjiang in CBM, which was distantly related to Kuriyama virus and Mukawa virus strains, with low nt identities of 75.8–77.4% for L segment, 67.6–68.7% for M segment M, and 69.8– 70.8 % for S segment (Fig. 5B, Extended data Table E9, 10) (40).

STPV and OTPV NE strains found in this study were clustered with STPV and OTPV strains discovered in *I. persulcatus* ticks from Karelia in Russia, with nt identities of 98.0–99.2% and 98.6–99.1%, respectively (Fig. 5C, Extended data Table E11, 12). The two viruses formed separate clades from each other, with nt identities of 56.4-57.3%, and were identified in paired in *I. persulcatus* ticks in six libraries from three collection sites, including Yichun in XXAM, and Tahe and Songling in DXAM (Fig. 5C, Extended data Table E11, 12).

#### 3.4.2.4 Rhabdoviridae

There were three novel rhabdoviruses identified in the study, namely, Tahe rhabdovirus 1 (THRV1), Tahe rhabdovirus 2 (THRV2), and Tahe rhabdovirus 3 (THRV3); they, together with Bole tick virus 2, Tacheng tick virus 3, Huangpi tick virus 3, and Wuhan tick virus 1, were genetically grouped into an unclassified Alphanemrhavirus-like group in the family *Rhabdoviridae* (Fig. 6A). A total of eight libraries of *H. japonica, H. concinna*, and D. *silvarum* ticks were detected THRV1, which formed close relationship with Manly virus, with nt identities of 30.4–30.6% and RdRp aa identities of 70.2–70.7%, respectively (Fig. 6B, Extended data Table E13). However, THRV1 were divided into two different clades: one clade was identified from libraries of Ji’an, Dunhua, Shulan, and Yichun in CBM and XXAM, while the other clade found in libraries from Tahe and Songling sited in DXAM, with nt identities of 81.4% and RdRp aa identities of 93.6–93.7% (Fig. 6B, Extended data Table E13). Compared with THRV1, THRV2 and THRV3 were only identified in the *I. persulcatus* tick from Tahe in DXAM, with nt identities of 44.9–49.3% with THRV1 and 45.9–46.1% with each other (Fig. 6B, Extended data Table E13).

**Figure 6.**
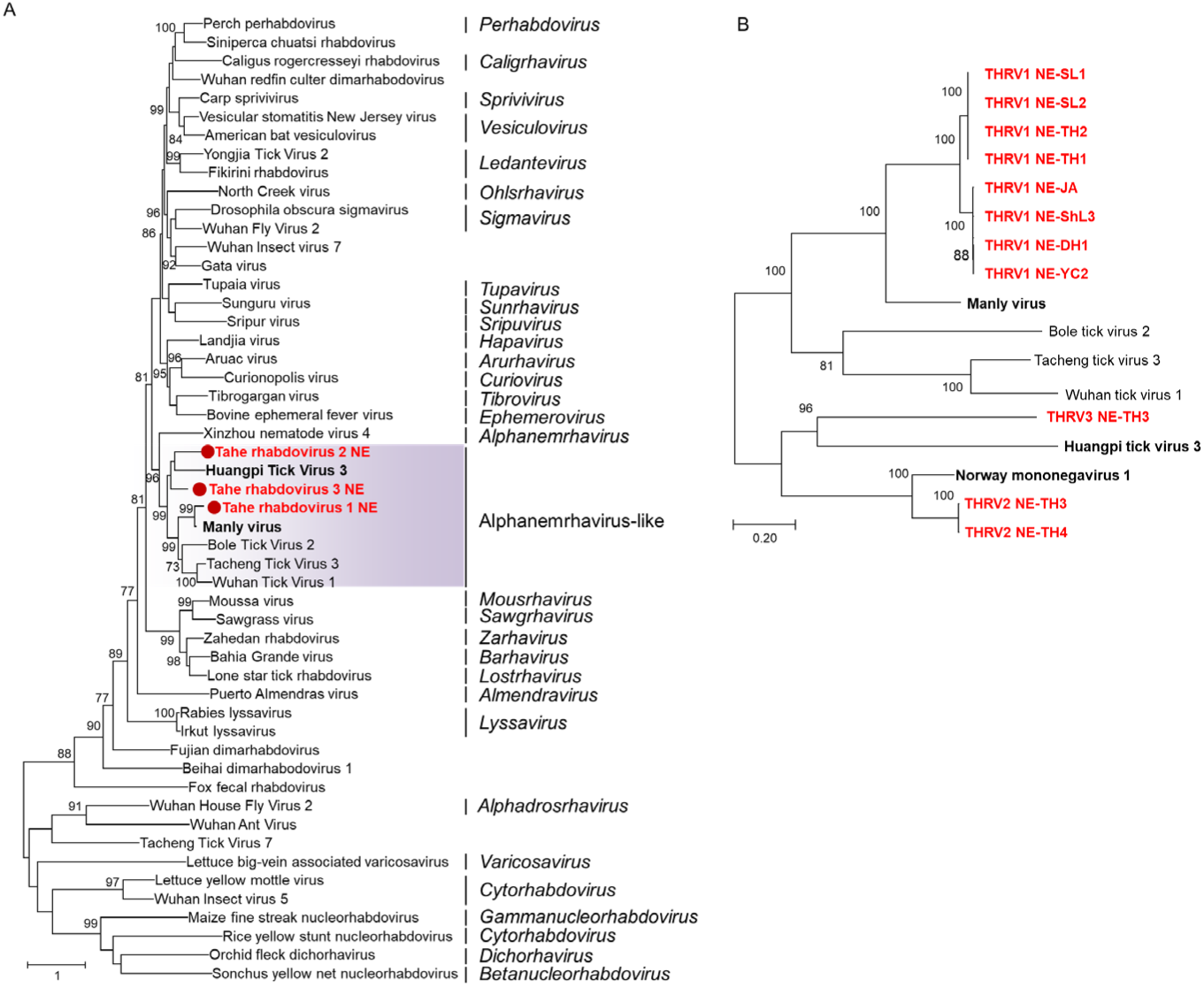
Phylogenetic analyses of rhabdoviruses. Phylogenetic trees constructed based on the RdRp sequences of representative viruses in the family *Rhabdoviridae* (A), the RdRp sequences of novel identified Tahe rhabdoviruses (B). All the viruses obtained in ticks here were highlighted in red, and the closest referenced viruses were also highlighted in bold font. Abbreviations: THRV1, Tahe rhabdovirus 1; THRV2, Tahe rhabdovirus 2; THRV3, Tahe rhabdovirus 3; OTPV, Onega tick phlebovirus; NWMV1: Norway mononegavirus 1; MLV: Manly virus; HPTV3: Huangpi tick virus 3; BLTV2: Bole tick virus 2; Tacheng tick virus 3: TCTV3; WHTV1: Wuhan tick virus 1; XZDV1: Xinzhou dimarhabdovirus virus 1. The accession numbers of the viral sequences used in the trees are shown in Supplementary Table S3 and S4.

#### 3.4.2.5 Chuviridae

Nuomin virus (NUMV) and Yichun mivirus (YCMV) identified here fell within the genera *Mivirus* and *Nigecruvirus in the family Chuviridae*, respectively (Fig. 7A). NUMV NE strains were detected in all the ten *I. persulcatus* libraries in the three regions in NE China, while no libraries from other tick species were identified NUMV positive, indicating the wide distribution and specificity of host tick species of the virus. The viral strains of NUMV found in this study were clustered with other NUMV strains discovered in humans from NE China and Lesnoe mivirus isolated from the *I. persulcatus* ticks in Russia, showing nucleotide identities of 96.6– 99.5% with each other (Extended data Table E14).

**Figure 7.**
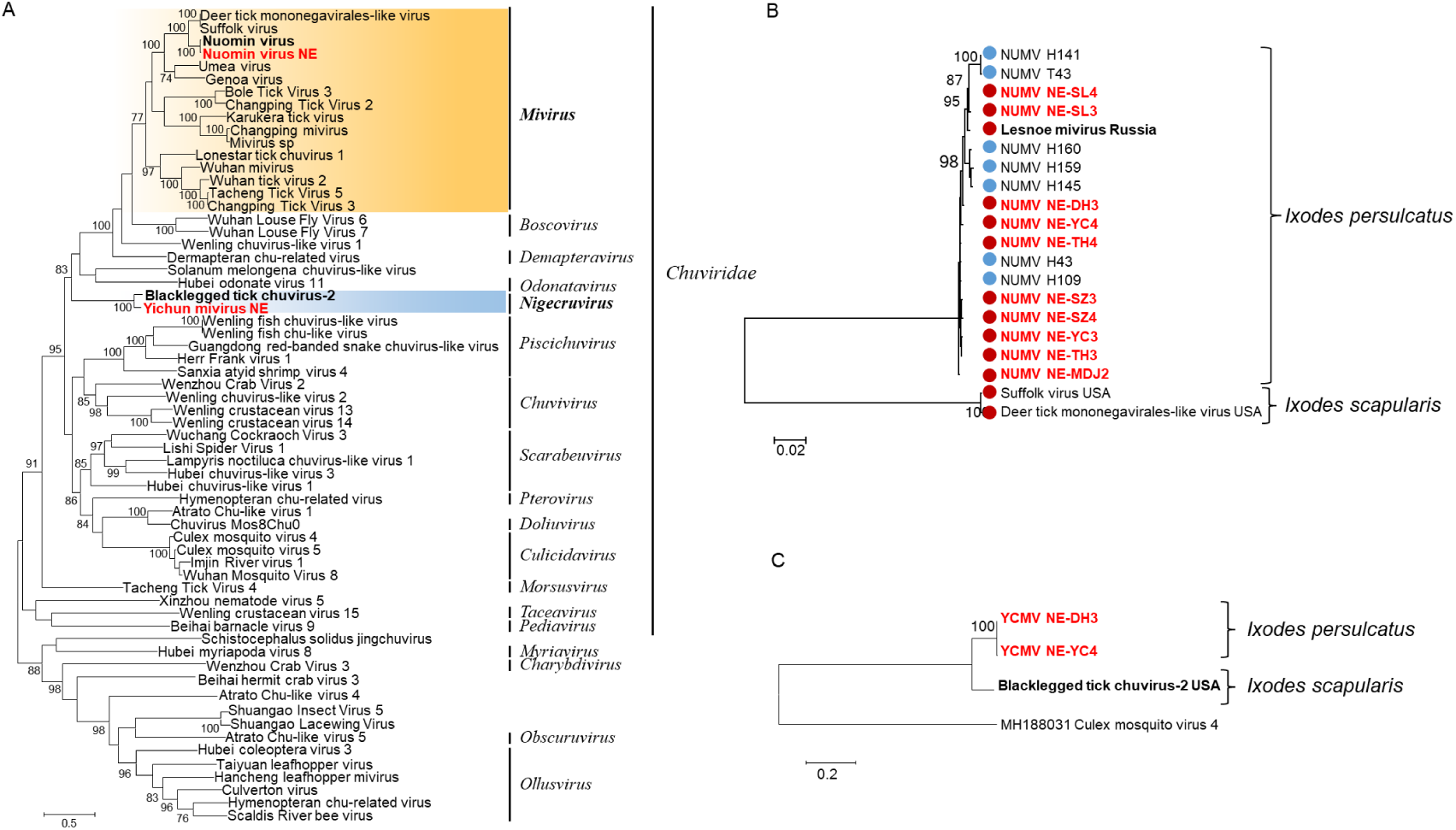
Phylogenetic analyses of chuviruses. Phylogenetic trees were constructed based on the RdRp sequences of representative viruses in the family *Chuviridae* (A), the RdRp sequences of NUMV (B) and JLCV (C). All the viruses obtained in ticks here were highlighted in red, and the closest referenced viruses were also highlighted in bold font. In panel B, NUMV viral strains discovered from ticks were marked with red-filled circles, while strains found in tick-bite patients were marked with blue-filled circles. Abbreviations: NUMV, Nuomin virus; YCMV, Yichun mivirus. The accession numbers of the viral sequences used in the trees are shown in Supplementary Table S3 and S4.

YCMV was a novel virus identified in this study, and clustered together with Blacklegged tick chuvirus 2 found in *I. scapularis* ticks from USA, with nt identities of 68.5%. YCMV was only detected in the *I. persulcatus* ticks from Dunhua and Yichun in CBM and XXAM (Fig. 7C), and shared high nt identity of 99.2%.

#### 3.4.2.6 Partitiviridae

The Jilin partiti-like virus 1 (JPLV1) identified here fell within an unclassified clade, provisionally designated as Deltapartitivirus-like group (Fig. 8A). JPLV1 NE strains were clustered with JPLV1 strains JL/QG 1, JL/QG 2, and JL/QG 4 isolated from *I. persulcatus* ticks in Jilin Province, NE China, with the nt identity of more than 99%, and JPLV1 was closely related to Norway partiti-like virus 1 strains with high identities of nt 91.3–91.9% and aa 94.1–94.3% (Extended data Table E15). Seven *I. persulcatus* libraries from all the three regions were identified JPLV1 positive, with nt identities of more than 99% to each other (Fig. 8B, Extended data Table E15).

**Figure 7.**
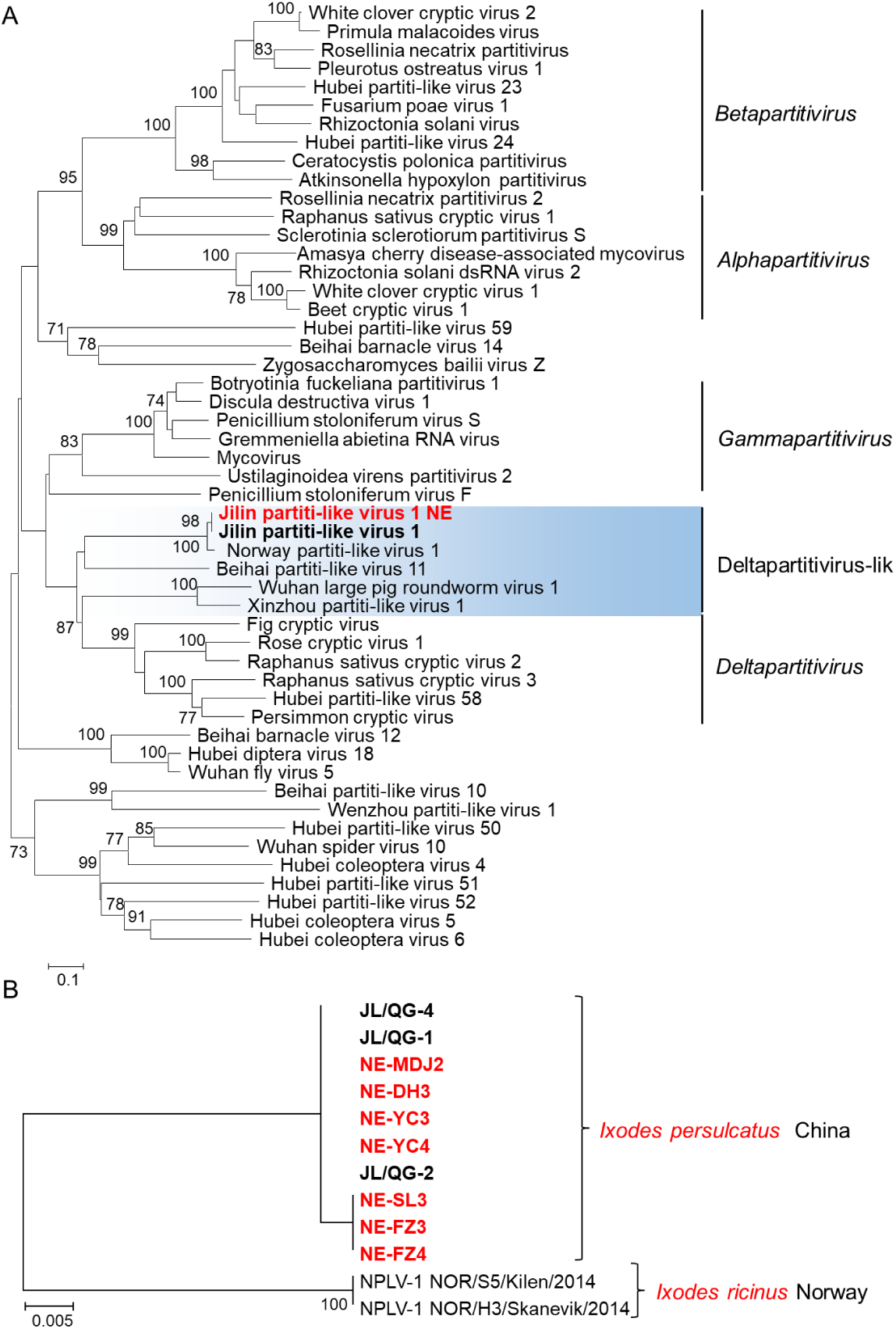
Phylogenetic analyses of partitiviruses. Phylogenetic trees constructed based on the RdRp protein sequences of representative viruses in the family *Partitiviridae* (A), the RdRp protein sequences of JPLV1 (B). All the viruses obtained in ticks here were highlighted in red, and the closest referenced viruses were also highlighted in bold font. Abbreviations: JPLV1, Jilin partiti-like virus 1; NPLV1, Norway partiti-like virus 1. The accession numbers of the viral sequences used in the trees are shown in Supplementary table S3 and S4.

**Figure 8.**
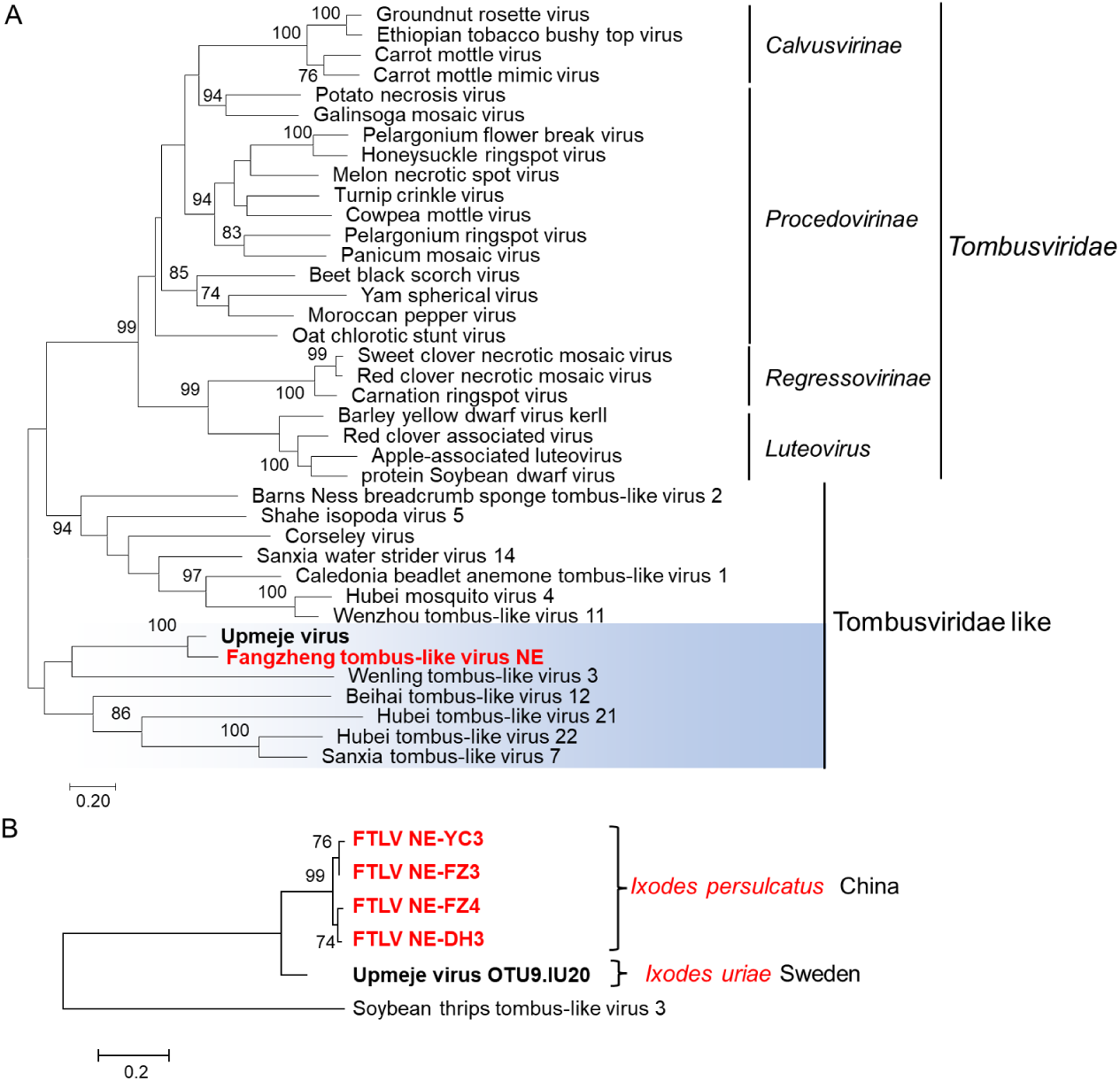
Phylogenetic analyses of tombusviruses. Phylogenetic trees were constructed based on the RdRp sequences of representative viruses in the family *Tombusviridae* (A), the RdRp sequences of FTLV (B). All the viruses obtained in ticks here were highlighted in red, and the closest referenced viruses were also highlighted in bold font. Abbreviations: FTLV, Fangzheng tombus-like virus. The accession numbers of the viral sequences used in the trees are shown in Supplementary Table S3 and S4.

#### 3.4.2.7 Tombusviridae

The newly discovered Fangzheng tombus-like virus (FTLV) fell within the Tombusviridae-like group in the family *Tombusviridae* (Fig. 8A). FTLV NE strains were only found in *I. persulcatus* tick from Yichun, Fangzheng, and Dunhua in XXAM and CBM (Fig. 8B), sharing a close relationship (identity: nt 73.4–73.6% and aa 80.6–81.3%) to Upmeje virus strain OTU9.IU20 that has been identified in *Ixodes uriae* in Sweden (Extended data Table E16) (41).

#### 3.4.2.8 Solemoviridae

*Ixodes scapularis* associated virus 1 (ISAV1), Xinjiang tick associated virus 1 (XTAV1), and Jilin luteo-like virus 2 (JLLV2) fell within the Sobemo-like virus group associated with the family *Solemoviridae* (Fig. 9A). In the aa RdRp phylogenetic tree, ISAV1-NE was clustered with Norway luteo-like virus 3 identified in *Ixodes Ricinus* ticks in Norway and *Ixodes scapularis* associated virus 1 identified in *I. scapularis* ticks in USA, with nt identities of 81.5–85.7% (Fig. 9B, Extended data Table E17). All the six ISAV1 NE strains were found in the *I. persulcatus* ticks from all the three regions in NE China, with nt identities of 98.1–100% with each other (Extended data Table E17).

**Figure 9.**
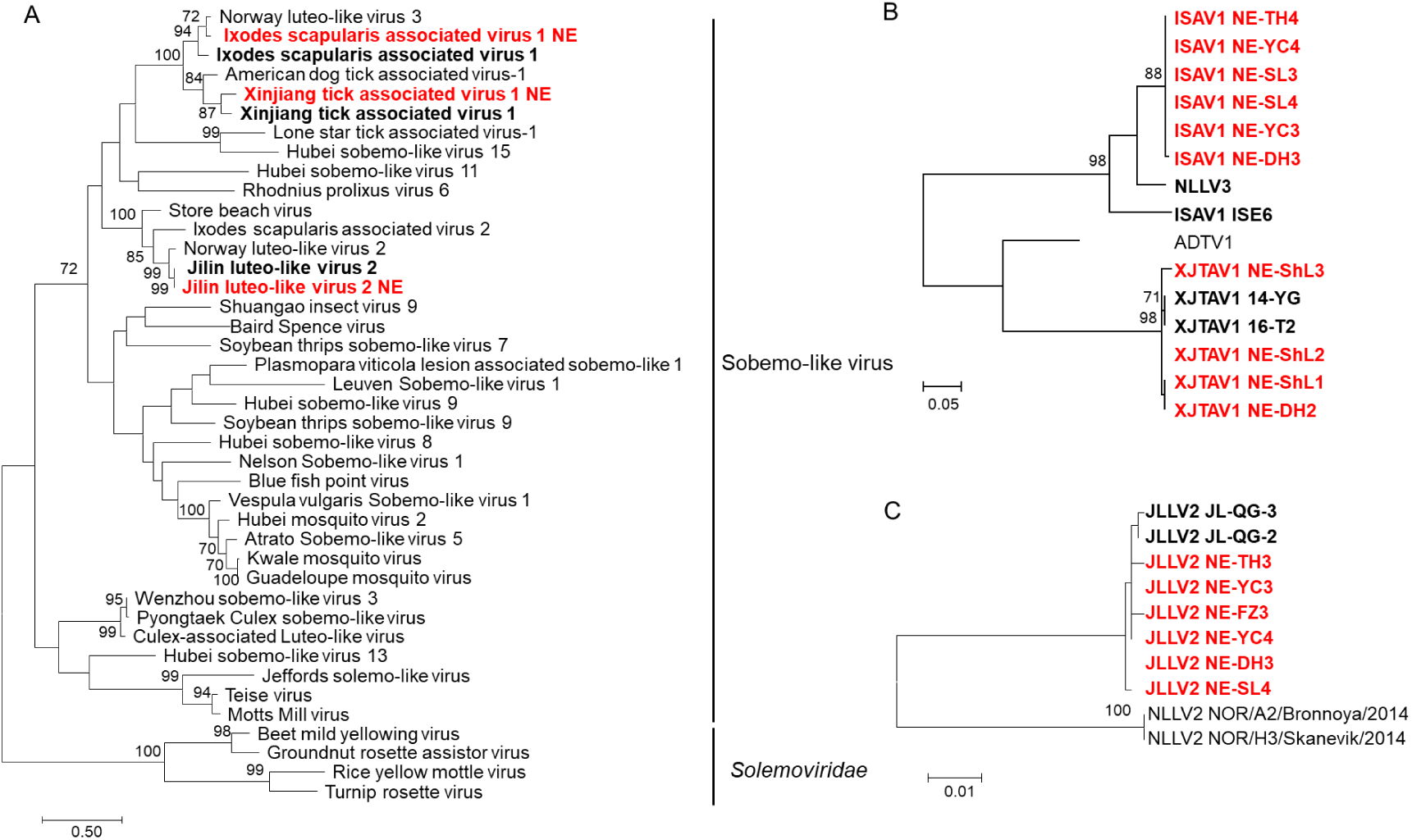
Phylogenetic analyses of Solemoviruses. Phylogenetic trees were constructed based on the RdRp sequences of representative viruses in the family *Solemoviridae* (A), the RdRp sequence of ISAV1 and XTAV1 (B), the RdRp sequence of JLLV2 (C). All the viruses obtained in ticks here were highlighted in red, and the closest referenced viruses were also highlighted in bold font. Abbreviations: ISAV1, *Ixodes scapularis* associated virus 1; XTAV1, Xinjiang tick associated virus 1; NLLV3, Norway luteo-like virus 3; ADTV, American dog tick associated virus-1; JLLV2, Jilin luteo-like virus 2; NLLV2, Norway luteo-like virus 2. The accession numbers of the viral sequences used in the trees are shown in Supplementary table S3 and S4.

XTAV1 NE strains formed a different clade from ISAV1, with nt identities of 61.9–62.8% (Fig. 9B, Extended data Table E17), and were detected in all the four *D. silvarum* tick pools collected from Shulan and Dunhua sited in CBM. The virus strains were clustered together with XJTAV1 strains 14-YG and 16-T2 identified in *Dermacentor nuttalli* in Xinjiang Uygur Autonomous Region, with nt identities of 95.9–96.2% (Extended data Table E17).

JLLV2 showed a close relationship with Norway luteo-like virus 2 identified in *Ixodes ricinus* ticks in Norway, with nt identities of 87.7–88.2% and aa identities of 91.3–91.7% (Fig. 9C, Extended data Table E18). JLLV2 NE strains were detected in the *I. persulcatus* ticks from six libraries from all the three regions, and clustered with JLLV2 strains in *I. persulcatus* from Jilin and shared the closest relationship with nt identity of more than 98% (Extended data Table E18).

#### 3.4.2.9 Unclassified

*Ixodes scapularis* associated virus 3 (ISAV3) NE strains identified in this study showed close relationship with ISAV3 and ISAV4 in *Ixodes scapularis* ticks from USA, but formed a different clade, with nt identities of 86.7–88.2% (Fig. 10, Extended data Table E19)(42, 43). Five libraries of *I. persulcatus* ticks from Tahe, Songling, and Yichun in DXAM and XXAM were detected ISAV3-positive, with nt identities of 93.7–99.8% (Fig. 10, Extended data Table E19).

**Figure 10.**
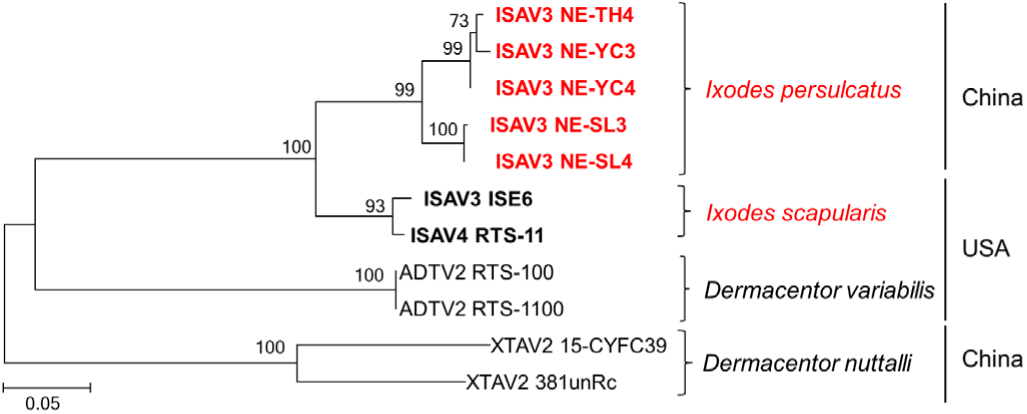
Phylogenetic trees constructed based on the genome sequences of ISAV3 and representative viruses. All the viruses obtained in ticks here were highlighted in red, and the closest referenced viruses were highlighted in bold font. The tick host and collection country of the virus strain were also marked. ISAV3, Ixodes scapularis associated virus 3; ISAV4, Ixodes scapularis associated virus 4; XTAV2, Xinjiang tick associated virus 2; ADTV, ISAV3, American dog tick associated virus 2. The accession numbers of the viral sequences used in the trees are shown in Supplementary Table S3 and S4.

## Discussion

We show the metagenomic description of the RNA viruses present in ticks in NE China. Analysis of the transcriptomes of *I. persulcatus, D. silvarum, H. conicinna*, and *H. japonica* ticks demonstrated that these ticks harbored a wide diversity of RNA viruses, belonging to at least 8 families of *Flaviviridae, Nairoviridae, Phenuiviridae, Rhabdoviridae, Chuviridae, Partitiviridae, Tombusviridae*, and *Solemoviridae*. Abundant viruses of the families *Flaviviridae, Nairoviridae, Phenuiviridae*, and *Chuviridae* detected in the ticks were consistent with those of previous studies, suggesting that viruses of these families have a wide geographical distribution (4, 35-37, 43-49).

Previous metagenomic analysis of ticks in Heilongjiang has revealed viral contigs annotated to South Bay virus (SBV), blacklegged tick phlebovirus (BTPV), deer tick Mononegavirales-like virus (DTMV), and Jingmen tick virus (JMTV) (21). Due to lack of whole genome sequence of these viruses, it is difficult to accurately define their classification [21]. In this study, we obtained the complete genome of 16 viral species, including 3 flaviviruses (ALSV, TBEV and BLTV-4), 4 nairoviruses (SGLV, JANV, BJNV, and YCNV), 4 phleboviruses (MV, KV, STPV, and OPTV), 3 rhabdoviruses (THRV-1, 2, and 3), and 2 chuviruses (NUMV and YCMV), showing an extensive diversity of RNA viruses in ticks in northeastern China.

There were significantly differences in tick viromes among CBM, XXAM, and DXAM. In this study, ALSV, TBEV, and THRV2-3 were only detected in DXAL, while MJPV, YCNV, YCMV, and FTLV were identified in CBM and XXAL. Nairovirus SGLV was detected in DXAM, while its closely related virus JANV was detected in CBM and XXAM; Rhabdovirus THRV1 tended to evolve into two genotypes: one was distributed in DXAM, the other distributed in CBM and XXAM. Additionally, the genetic differences of TBEV also supports the occurrence of geographical barriers of tick-borne viruses in northeastern China.

Segmented flaviviruses have recently been reported as emerging tick-borne viruses, with a wide distribution in Asia, Africa, Europe, Central America, and South America (44, 45). Of them, Jingmen tick virus (JMTV) and ALSV are associated with the febrile illness in tick-bitten patients in northeastern China. JMTV has been found in various vertebrates, including cattle, sheep, rodents, and non-human primate, showing that the virus cocirculates between ticks and mammals (44); JMTV and ALSV have also been detected in mosquitoes (14, 50); however, the transmission modes of these viruses remain to be investigated. In this study, only *I. persulcatus* tick was tested ALSV-positive in DXAM, where ALSV patients have been recently found (14), suggesting potential public health risk of ALSV infection and limited distribution of the emerging tick-borne virus in northeastern China. The pathogenicity of segmented flaviviruses in humans and animals needs to be further verified through animal infection models.

Interestingly, co-feeding transmission might affect the viromes in ticks collected from animals. In this study, two viral species, THRV1 and MKWV, identified in *D. silvarum* collected from cattle (Shulan), were mainly found in *H. japonica* and *H. concinna*, and *I. persulcatus* ticks, respectively. However, the two viruses were not detected in the questing *D. silvarum* ticks in Dunhua adjacent to Shulan. As cattle can act as hosts of more than two tick species, such as *D. silvarum, H. japonica, H. concinna*, and *I. persulcatus* ticks, and co-feeding transmission may be an efficiency way of viral transmission in ticks (51-53), we concluded that *D. silvarum* collected from cattle here may contract the viruses from *Haemaphysalis sp*. or *I. persulcatus* ticks by co-feeding transmission, but not the actual vector of these viruses. Thus, it is suggested to investigate tick virome diversity using questing ticks instead of ticks collected from animals. Moreover, further studies should be focus on the vector competence of ticks for the transmission of the identified viruses, which may further confirm the roles of different tick species for virus transmission.

The Bunyavirales order includes important human pathogens, such as Crimean-Congo hemorrhagic fever virus in the *Nairoviridae* family, and SFTSV and Valley fever virus in the *Phenuiviridae* family, whose genome are negative single-stranded RNA of small (S), medium (M), and large (L) segments, encoding structural nucleoprotein (NP), glycoprotein precursor (GPC), and RNA-dependent RNA polymerase (L) proteins, respectively. Recently, several bi-segmented viruses without M gene have been found in nairoviruses (e.g., Gakugsa tick virus and Norway nairovirus 1) and phleboviruses (e.g., Tacheng tick virus 2) (46-49). We identified four viruses including YCNV and BJNV within the family *Nairoviridae*, and STPV together with OTPV in the family *Phenuiviridae*. However, we did not find the M gene encoding the glycoproteins. The possible reason may be the insufficient homology between these viruses and reference viruses (16, 18).

We also found several viral species closely related to plant viruses, including JPLV1 in the family *Partitiviridae*, FTLV of with family *Tombusviridae*, and ISAV1, XTAV1, and JLLV2 in family *Solemoviridae*. Compared with viruses in family *Flaviviridae, Nairoviridae*, and *Phenuiviridae*, plant-related viruses identified here may possess low pathogenic to humans or mammals.

There are some limitations to the present study. Although some tick-borne viruses associated with diseases in humans or mammals, including TBEV, ALSV, SGLV, BJNV, and NUMV, were detected in this study, some other pathogenic virus, such as severe fever with thrombocytopenia syndrome virus (SFTSV) (54, 55), Lymphocytic choriomeningitis virus (LCMV) (56), Nairobi sheep disease virus (NSDV) (19), and Jingmen tick virus (JMTV) (13) identified in previous studies, were not detected here, indicating that larger tick samples and wider sampling sites are necessary to characterize the tick-borne viruses in NE China. Some novel viruses had close relationship with TBVs of public health significance. For example, JANV and YCNV were genetically related to SGLV and BJNV, respectively, which have been shown to be associated with febrile diseases, indicating the potentially pathogenic to humans and animals of these novel viruses. As in the family *Phenuiviridae*, however, although MKWV does not detected in tick-bitten patients, it can grow in a human-derived cell line (Huh-7) and mice, and its NSs can function the anti-innate immune responses (40), suggesting the potential public health significance of MKWV and its closely related virus (MJPV), and the necessity of epidemiology studies on tick-bitten patients, livestock, and even wild animals.

The viruses identified in this study revealed an extensive diversity of RNA viruses in ticks, and provided insights into the geographical distribution and transmission of TBVs in northeastern China, highlighting the necessity of surveillance and laying a foundation for further study on tick-borne viruses. Moreover, considering the potential pathogenicity of some novel viruses, epidemiology studies of tick-bitten patients and animals, establishment of animal infection models, and vector competence verification of ticks on these viruses should be done to further evaluate their public health significance.

## Acknowledgments

This study was supported by the National Natural Science Foundation of China (82002165 and 32072887); Outstanding Young Scholars Cultivating Plan of the First Hospital of Jilin University (2021-YQ-01); the Pearl River Talent Plan in Guangdong Province of China (2019CX01N111); and Medical and Health Talent Special Project in Jilin Province of China (JLSWSRCZX2021-002)

## About the Author

Ms. Liu is a master student in the School of Life Science of Jilin University. Her research interest is tick-borne infectious disease epidemiology.

## Data availability

All sequence reads have been deposited in the Short Read Archive BioProject XXX. Viral genomes have been submitted to GenBank with Accession Nos XXX–XXX, showed in Supplementary Table S4

## Supplementary data

Supplementary data is available online.

## Author contributions

Z.W. and Q.L. designed the research. Z.W., Z.L., F.W., L.L., Y.C., Q.Y., Z.H., and D.L. collected the samples. Z.L., W.X., Y.Y., X.L., Y. Z, L.S., Q.L., L.Z., Z.W., and H.Z. performed the experiments. Z.W. and Z.L. analyzed the data. Z.W. and Z.L. wrote the original manuscript. Z.W. and Q.L. reviewed and edited the manuscript. All authors gave final approval for publication.

