## Supplementary material for "Extensive diversity of RNA viruses in ticks revealed by metagenomics in northeastern China": Extended data.docx

Table S1. Nucleotide sequence identities of S1 (upper right) and S2 (lower left) segments of ALSV calculated using MegAlign program available within DNAstar V7.1.*

|  | ALSV TH4 | ALSV H3 | ALSV HLJ1 | ALSV HLJ2 | ALSV Miass527 | ALSV Miass502 | ALSV Miass519 | ALSV Miass506 | ALSV Kuutsalo-23 | ALSV Haapasaari-18 | TKCV IM-OI70 | XJTV1 XJO381 | YGTV XJ-YGTV-1 | HLJTV HLJ41 | JMTV HLJ41 | GXTV GX46 |
| --- | --- | --- | --- | --- | --- | --- | --- | --- | --- | --- | --- | --- | --- | --- | --- | --- |
| ALSV TH4 | *** | 97 | 95.8 | 95.8 | 89.6 | 89.4 | 88.9 | 88.9 | 89.2 | 89 | 73.6 | 72.8 | 72.1 | 71.8 | 71.9 | 71.8 |
| ALSV H3 | 98.2 | *** | 95.5 | 95.5 | 89.6 | 89.4 | 89 | 89 | 89.6 | 89.5 | 74.2 | 73.1 | 72.3 | 72.2 | 72 | 72.2 |
| ALSV HLJ1 | 94.2 | 94.8 | *** | 100 | 89.3 | 89.2 | 88 | 88 | 89.4 | 89.2 | 73.5 | 72.9 | 71.7 | 71.2 | 71.2 | 71.2 |
| ALSV HLJ2 | 94.2 | 94.8 | 100 | *** | 89.3 | 89.2 | 88 | 88 | 89.4 | 89.2 | 73.5 | 72.9 | 71.7 | 71.2 | 71.2 | 71.2 |
| ALSV Miass527 | 93.9 | 94.2 | 96.7 | 96.7 | *** | 99.5 | 90.7 | 90.7 | 89.4 | 88.8 | 73.8 | 72.2 | 71.8 | 71.1 | 71 | 71.1 |
| ALSV Miass502 | 93.8 | 94.1 | 96.5 | 96.5 | 99.5 | *** | 90.5 | 90.5 | 89.2 | 88.7 | 73.6 | 72.1 | 71.8 | 71 | 70.9 | 71 |
| ALSV Miass519 | 92.1 | 91.8 | 92.2 | 92.2 | 92.4 | 92.3 | *** | 100 | 89.1 | 88.8 | 73.8 | 72.3 | 71.6 | 71.9 | 71.6 | 71.9 |
| ALSV Miass506 | 92.1 | 91.8 | 92.2 | 92.2 | 92.4 | 92.3 | 100 | *** | 89.1 | 88.8 | 73.8 | 72.3 | 71.6 | 71.9 | 71.6 | 71.9 |
| ALSV Kuutsalo-23 | 91.3 | 91.1 | 91.2 | 91.2 | 92 | 91.9 | 93.6 | 93.6 | *** | 95.8 | 74.2 | 71.8 | 72.1 | 71.3 | 71.1 | 71.3 |
| ALSV Haapasaari-18 | 91.3 | 91 | 90.8 | 90.8 | 91.7 | 91.6 | 93.9 | 93.9 | 98.4 | *** | 74.1 | 71.3 | 72 | 71.7 | 71.1 | 71.7 |
| TKCV IM-OI70 | 69.1 | 68.8 | 68.9 | 68.9 | 69.1 | 69.1 | 67.4 | 67.4 | 68.5 | 68.4 | *** | 72.2 | 72.1 | 71.6 | 71.1 | 71.6 |
| XJTV1 XJO381 | 64.9 | 64.8 | 64.9 | 64.9 | 65.7 | 65.7 | 65.1 | 65.1 | 65.1 | 65 | 62.6 | *** | 77.2 | 70.3 | 69.8 | 70.3 |
| YGTV XJ-YGTV-1 | 64.6 | 64.5 | 65.2 | 65.2 | 65.6 | 65.4 | 65.6 | 65.6 | 66 | 66 | 64.6 | 76.2 | *** | 69.7 | 71 | 69.7 |
| HLJTV HLJ41 | 59.7 | 59.6 | 60.4 | 60.4 | 59.8 | 59.6 | 59.2 | 59.2 | 59.8 | 59.7 | 58.7 | 58.6 | 58.6 | *** | 93 | 100 |
| JMTV HLJ41 | 58.8 | 58.7 | 59.8 | 59.8 | 58.8 | 58.8 | 58.7 | 58.7 | 59.6 | 59.6 | 59 | 57.8 | 57.5 | 94.3 | *** | 93 |
| GXTV GX46 | 59.7 | 59.6 | 60.4 | 60.4 | 59.8 | 59.6 | 59.2 | 59.2 | 59.8 | 59.7 | 58.7 | 58.6 | 58.6 | 100 | 94.3 | *** |

* Abbreviations: ALSV, Alongshan virus; TKCV, Takachi virus; XJTV, Xinjiang tick virus; YGTV, Yanggou tick virus; HLJTV, Heilongjiang tick virus; JMTV, Jingmen tick virus; GXTV, Guangxi tick virus.

Table S2. Nucleotide sequence identities of S3 (upper right) and S4 (lower left) segments of ALSV calculated using MegAlign program available within DNAstar V7.1.*

|  | ALSV TH4 | ALSV H3 | ALSV HLJ1 | ALSV HLJ2 | ALSV Miass527 | ALSV Miass502 | ALSV Miass519 | ALSV Miass506 | ALSV Kuutsalo-23 | ALSV Haapasaari-18 | TKCV IM-OI70 | XJTV1 XJO381 | YGTV XJ-YGTV-1 | HLJTV HLJ41 | JMTV HLJ41 | GXTV GX46 |
| --- | --- | --- | --- | --- | --- | --- | --- | --- | --- | --- | --- | --- | --- | --- | --- | --- |
| ALSV TH4 | *** | 97.8 | 93.2 | 93.2 | 91 | 91.1 | 90.9 | 90.9 | 90 | 90.5 | 75.6 | 73.2 | 72.8 | 72.3 | 72 | 72.3 |
| ALSV H3 | 98.5 | *** | 93.1 | 93.1 | 90.1 | 90.2 | 90.5 | 90.5 | 90.3 | 90.3 | 75.4 | 73.4 | 72.9 | 72.2 | 72.2 | 72.2 |
| ALSV HLJ1 | 97.3 | 97 | *** | 100 | 90.3 | 90.5 | 91.1 | 91.1 | 90.2 | 90.3 | 76 | 73.4 | 73.5 | 72.2 | 72.1 | 72.2 |
| ALSV HLJ2 | 97.3 | 97 | 100 | *** | 90.3 | 90.5 | 91.1 | 91.1 | 90.2 | 90.3 | 76 | 73.4 | 73.5 | 72.2 | 72.1 | 72.2 |
| ALSV Miass527 | 89.6 | 89.6 | 89.5 | 89.5 | *** | 99.8 | 96.7 | 96.7 | 90.1 | 89.3 | 76 | 73.6 | 73.9 | 72.1 | 72 | 72.1 |
| ALSV Miass502 | 89.4 | 89.4 | 89.2 | 89.2 | 99.5 | *** | 96.9 | 96.9 | 90.1 | 89.4 | 76 | 73.5 | 74 | 72.1 | 72 | 72.1 |
| ALSV Miass519 | 89.5 | 89.5 | 89.1 | 89.1 | 98.5 | 98.1 | *** | 100 | 90.3 | 89.4 | 76 | 73.2 | 73.7 | 72.4 | 72.4 | 72.4 |
| ALSV Miass506 | 89.5 | 89.5 | 89.1 | 89.1 | 98.5 | 98.2 | 100 | *** | 90.3 | 89.4 | 76 | 73.2 | 73.7 | 72.4 | 72.4 | 72.4 |
| ALSV Kuutsalo-23 | 90.1 | 90.1 | 89.6 | 89.6 | 90.1 | 89.8 | 90.1 | 90.1 | *** | 94.4 | 75.6 | 72.8 | 73.2 | 71.2 | 70.9 | 71.2 |
| ALSV Haapasaari-18 | 89.6 | 89.7 | 89 | 89 | 90.2 | 89.9 | 90 | 90 | 94 | *** | 75.8 | 72.6 | 73.7 | 71.9 | 71.6 | 71.9 |
| TKCV IM-OI70 | 72.8 | 73 | 72.9 | 72.9 | 72.6 | 72.3 | 73 | 73 | 72.6 | 72.8 | *** | 73 | 73.8 | 71.7 | 70.9 | 71.7 |
| XJTV1 XJO381 | 69.3 | 69.4 | 68.8 | 68.8 | 69.3 | 68.9 | 69.3 | 69.2 | 70 | 68.8 | 68.4 | *** | 77.6 | 70.8 | 70.6 | 70.8 |
| YGTV XJ-YGTV-1 | 68.4 | 68.3 | 68.5 | 68.5 | 68.1 | 67.7 | 68.1 | 68 | 68.6 | 68.1 | 68.2 | 76.1 | *** | 70.2 | 69.7 | 70.2 |
| HLJTV HLJ41 | 64.9 | 64.9 | 64.8 | 64.8 | 65.4 | 65.1 | 65 | 64.9 | 65.1 | 65.3 | 65.9 | 63.1 | 62.2 | *** | 94.7 | 100 |
| JMTV HLJ41 | 64.5 | 64.5 | 64.4 | 64.4 | 65.2 | 64.9 | 65.1 | 65.1 | 65.5 | 65.4 | 65.2 | 62.8 | 62 | 92.7 | *** | 94.7 |
| GXTV GX46 | 64.9 | 64.9 | 64.8 | 64.8 | 65.4 | 65.1 | 65 | 64.9 | 65.1 | 65.3 | 65.9 | 63.1 | 62.2 | 100 | 92.7 | *** |

* Abbreviations: ALSV, Alongshan virus; TKCV, Takachi virus; XJTV, Xinjiang tick virus; YGTV, Yanggou tick virus; HLJTV, Heilongjiang tick virus; JMTV, Jingmen tick virus; GXTV, Guangxi tick virus.

Table S3. Nucleotide sequence identities of TBEV calculated using MegAlign program available within DNAstar V7.1.*

|  | NE-TH3 | NE-TH4 | SL-TH4 | DXAL_T83 | HLB-T74 | JL_Jiaohe | JL-T75 | Senzhang | JLCB11-08 | JLCB11-35 | JLCB11-40 | MDJ01 | MDJ-03 | MDJ-02 |
| --- | --- | --- | --- | --- | --- | --- | --- | --- | --- | --- | --- | --- | --- | --- |
| NE-TH3 | *** | 93.5 | 97.7 | 93.5 | 93.5 | 94.3 | 93.8 | 93.5 | 93.9 | 93.9 | 93.9 | 93.5 | 93.6 | 93.6 |
| NE-TH4 | *** | *** | 93.9 | 98 | 98 | 93.7 | 95 | 94.8 | 95 | 95 | 95 | 95 | 94.8 | 94.8 |
| SL-TH4 | *** | *** | *** | 93.8 | 93.8 | 94.6 | 94.2 | 93.9 | 94.1 | 94.2 | 94.2 | 93.9 | 94 | 94 |
| DXAL_T83 | *** | *** | *** | *** | 99.7 | 93.5 | 94.7 | 94.4 | 94.7 | 94.7 | 94.7 | 94.6 | 94.5 | 94.5 |
| HLB-T74 | *** | *** | *** | *** | *** | 93.5 | 94.8 | 94.4 | 94.7 | 94.8 | 94.7 | 94.6 | 94.5 | 94.6 |
| JL_Jiaohe | *** | *** | *** | *** | *** | *** | 94.1 | 93.9 | 93.9 | 93.9 | 94 | 93.9 | 93.9 | 94 |
| JL-T75 | *** | *** | *** | *** | *** | *** | *** | 98.1 | 98.3 | 98.3 | 98.3 | 98.1 | 98.1 | 98.1 |
| Senzhang | *** | *** | *** | *** | *** | *** | *** | *** | 98.1 | 98.1 | 98.2 | 98 | 99.6 | 99.6 |
| JLCB11-08 | *** | *** | *** | *** | *** | *** | *** | *** | *** | 99.9 | 99.3 | 97.9 | 98.1 | 98.1 |
| JLCB11-35 | *** | *** | *** | *** | *** | *** | *** | *** | *** | *** | 99.3 | 98 | 98.1 | 98.1 |
| JLCB11-40 | *** | *** | *** | *** | *** | *** | *** | *** | *** | *** | *** | 98 | 98.2 | 98.2 |
| MDJ01 | *** | *** | *** | *** | *** | *** | *** | *** | *** | *** | *** | *** | 98.2 | 98.2 |
| MDJ-03 | *** | *** | *** | *** | *** | *** | *** | *** | *** | *** | *** | *** | *** | 99.9 |
| MDJ-02 | *** | *** | *** | *** | *** | *** | *** | *** | *** | *** | *** | *** | *** | *** |

Table S4. Nucleotide sequence identities of complete genome (upper right) and amino acid sequence identities of RdRp (lower left) of BLTV4 calculated using MegAlign program available within DNAstar V7.1.

|  | ShL3 | DH2 | Iasi23 | Iasi21 | Iasi20 | Iasi50 | 17-L2 | bole4-xinjiang-JMN | GSC346flaviV | BLP-1 | Bangali/H.truncatum/2018 | Iftin/H.dromedarii/2018 | TTP-Pool-4 | Thailand tick flavivirus |
| --- | --- | --- | --- | --- | --- | --- | --- | --- | --- | --- | --- | --- | --- | --- |
| ShL3 | *** | 98.1 | 82.7 | 82.5 | 82.7 | 82.8 | 78.9 | 78.9 | 78.9 | 78.8 | 78.5 | 78.8 | 78 | 77.5 |
| DH2 | 98.5 | *** | 82.9 | 82.7 | 82.9 | 83 | 79 | 79.1 | 79.1 | 79 | 78.7 | 79 | 78.2 | 77.6 |
| Iasi23 | 95.4 | 95.4 | *** | 99.5 | 99.8 | 97.3 | 80.4 | 80.4 | 80.4 | 80.3 | 80.3 | 80.3 | 79.2 | 79.4 |
| Iasi21 | 95 | 95 | 99.6 | *** | 99.5 | 97 | 80.2 | 80.2 | 80.2 | 80.1 | 80.1 | 80.1 | 79 | 79.2 |
| Iasi20 | 95 | 95 | 99.6 | 99.2 | *** | 97.3 | 80.3 | 80.4 | 80.4 | 80.3 | 80.3 | 80.3 | 79.2 | 79.4 |
| Iasi50 | 94.6 | 94.6 | 99.2 | 99.6 | 98.8 | *** | 80.3 | 80.3 | 80.3 | 80.2 | 80.2 | 80.2 | 79.2 | 79.3 |
| 17-L2 | 95.4 | 95.8 | 93.8 | 93.5 | 94.2 | 93.1 | *** | 98.9 | 98.5 | 98.6 | 88.1 | 91.5 | 79.9 | 80.4 |
| bole4-xinjiang-JMN | 95.8 | 96.2 | 94.2 | 93.8 | 94.6 | 93.5 | 99.6 | *** | 98.5 | 98.8 | 88.1 | 91.5 | 80 | 80.5 |
| GSC346flaviV | 96.2 | 96.2 | 94.2 | 93.8 | 93.8 | 93.5 | 98.8 | 99.2 | *** | 98.2 | 88 | 91.3 | 79.9 | 80.4 |
| BLP-1 | 96.2 | 96.5 | 94.6 | 94.2 | 94.2 | 93.8 | 99.2 | 99.6 | 99.6 | *** | 88.1 | 91.4 | 79.9 | 80.4 |
| Bangali/H.truncatum/2018 | 96.2 | 96.2 | 95.4 | 95 | 95 | 94.6 | 96.9 | 97.3 | 97.3 | 97.7 | *** | 92.6 | 84.5 | 83.7 |
| Iftin/H.dromedarii/2018 | 96.9 | 96.9 | 96.2 | 95.8 | 95.8 | 95.4 | 97.7 | 98.1 | 98.1 | 98.5 | 99.2 | *** | 81.7 | 81.4 |
| TTP-Pool-4 | 95 | 95 | 95.4 | 95 | 95 | 94.6 | 95 | 95.4 | 95.4 | 95.8 | 96.5 | 97.3 | *** | 89.7 |
| Thailand tick flavivirus | 95.8 | 95.8 | 95 | 95.4 | 94.6 | 95 | 95.8 | 96.2 | 96.2 | 96.5 | 96.5 | 97.3 | 98.5 | *** |

Table S5. Nucleotide sequence identities of L segment (upper right) and amino acid sequence identities of RdRp (lower left) of JANV and SGLV.

|  | JANV MDJ1 | JANV YC1 | JANV JA | JANV DH1 | SGLV TH3 | SGLV TH4 | SGLV YC585 | SGLV HLJ1202 | TCTV1 TC253 | SXTV2 SXO338nairoV | HNTV HNO321nairoV |
| --- | --- | --- | --- | --- | --- | --- | --- | --- | --- | --- | --- |
| JANV MDJ1 | *** | 99.2 | 99.1 | 98.4 | 73.3 | 73.4 | 73.4 | 73.1 | 61.6 | 63.2 | 64.6 |
| JANV YC1 | 99.7 | *** | 99.2 | 98.5 | 73.4 | 73.4 | 73.5 | 73.2 | 61.6 | 63.2 | 64.7 |
| JANV JA | 99.4 | 99.6 | *** | 98.7 | 73.3 | 73.4 | 73.4 | 73.1 | 61.5 | 63.2 | 64.6 |
| JANV DH1 | 99.6 | 99.7 | 99.6 | *** | 73.4 | 73.4 | 73.5 | 73.2 | 61.7 | 63.4 | 64.6 |
| SGLV TH3 | 85.1 | 85.2 | 85 | 85.1 | *** | 96.5 | 95.9 | 95.7 | 61.8 | 63.3 | 64 |
| SGLV TH4 | 85 | 85.1 | 84.9 | 85 | 99.4 | *** | 97 | 96.5 | 61.9 | 63.4 | 63.8 |
| SGLV YC585 | 84.8 | 84.9 | 84.7 | 84.8 | 99.1 | 99.2 | *** | 98.4 | 61.8 | 63.4 | 63.6 |
| SGLV HLJ1202 | 84.1 | 84.2 | 84 | 84.1 | 98.4 | 98.4 | 98.7 | *** | 61.7 | 63 | 63.4 |
| TCTV1 TC253 | 63.3 | 63.5 | 63.3 | 63.4 | 64 | 63.9 | 63.6 | 63.1 | *** | 62.3 | 61.7 |
| SXTV2 SXO338nairoV | 66 | 66.2 | 66 | 66.1 | 66.1 | 66.2 | 65.9 | 65.5 | 63.7 | *** | 66.4 |
| HNTV HNO321nairoV | 67.6 | 67.7 | 67.6 | 67.7 | 67.7 | 67.7 | 67.4 | 67 | 64.1 | 71.5 | *** |

* Abbreviations: JANV, Ji’an nariovirus; SGLV, Songling virus; TCTV1, Tacheng tick virus 1; SXTV2, Shanxi tick virus 2; HNTV, Henan tick virus.

Table S6. Nucleotide sequence identities of M segment (upper right) and S segment (lower left) of JANV and SGLV.

|  | JANV MDJ1 | JANV YC1 | JANV JA | JANV DH1 | SGLV TH3 | SGLV TH4 | SGLV YC585 | SGLV HLJ1202 | TCTV1 TC253 | SXTV2 SXO338nairoV | HNTV HNO321nairoV |
| --- | --- | --- | --- | --- | --- | --- | --- | --- | --- | --- | --- |
| JANV MDJ1 | *** | 99.3 | 99.7 | 99.2 | 71.2 | 71.6 | 71.1 | 71.1 | 54.6 | 57.5 | 60.3 |
| JANV YC1 | 99.1 | *** | 99.3 | 99.8 | 71.1 | 71.5 | 71 | 70.9 | 54.7 | 57.5 | 60 |
| JANV JA | 99 | 98.8 | *** | 99.5 | 71.2 | 71.5 | 71.1 | 71 | 54.6 | 57.5 | 60.2 |
| JANV DH1 | 97.8 | 97.4 | 97.5 | *** | 71.1 | 71.4 | 71 | 70.9 | 54.7 | 57.4 | 60 |
| SGLV TH3 | 71.3 | 71.4 | 71.1 | 71.7 | *** | 92.4 | 98.1 | 97.6 | 55.4 | 57.3 | 61.4 |
| SGLV TH4 | 71.1 | 71.1 | 71.1 | 71.5 | 99.1 | *** | 92.3 | 92.1 | 54.9 | 57.3 | 61.8 |
| SGLV YC585 | 71.3 | 71.3 | 71.1 | 71.7 | 99.2 | 99.1 | *** | 98.9 | 55.3 | 57.4 | 61.6 |
| SGLV HLJ1202 | 70.9 | 70.9 | 70.7 | 71 | 98.5 | 98.4 | 99 | *** | 55.1 | 57.1 | 61.5 |
| TCTV1 TC253 | 59.7 | 59.7 | 59.9 | 60.4 | 59.2 | 59.2 | 59 | 58.7 | *** | 56.1 | 56.9 |
| SXTV2 SXO338nairoV | 61.5 | 61.5 | 61.5 | 61.5 | 62.1 | 62.2 | 61.9 | 61.4 | 58 | *** | 59.4 |
| HNTV HNO321nairoV | 61.6 | 61.3 | 61.5 | 61.7 | 62.3 | 62.5 | 62.3 | 61.7 | 60.9 | 66.7 | *** |

* Abbreviations: JANV, Ji’an nariovirus; SGLV, Songling virus; TCTV1, Tacheng tick virus 1; SXTV2, Shanxi tick virus 2; HNTV, Henan tick virus.

Table S7. Nucleotide sequence identities of L segment (upper right) and amino acid sequence identities of RdRp (lower left) of YCNV and BJNV.

|  | YCNV YC4 | YCNV YC3 | YCNV FZ3 | BJNV SL4 | BJNV SL3 | BJNV TH3 | BJNV TH4 | BJNV YC4 | BJNV YC3 | BJNV DH3 | BJNV H1603 | BJNV H160 | BJNV H801 | BJNV H39 | BJNV H56 | BJNV H59 | GKTV | PTV | NWNV1 | GTV |
| --- | --- | --- | --- | --- | --- | --- | --- | --- | --- | --- | --- | --- | --- | --- | --- | --- | --- | --- | --- | --- |
| YCNV YC4 | *** | 98.6 | 98.7 | 83.1 | 83 | 83 | 83 | 83.1 | 83.1 | 83.1 | 82.9 | 82.9 | 82.9 | 82.9 | 82.5 | 82.5 | 82.7 | 77.8 | 77.7 | 77.5 |
| YCNV YC3 | 99.2 | *** | 99.4 | 83 | 83 | 83 | 83 | 83 | 83.1 | 83.1 | 82.8 | 82.8 | 82.8 | 82.8 | 82.5 | 82.5 | 82.7 | 77.7 | 77.7 | 77.4 |
| YCNV FZ3 | 99.2 | 99.6 | *** | 83.1 | 83 | 83 | 83 | 83.1 | 83.1 | 83.1 | 82.9 | 82.9 | 82.9 | 82.9 | 82.6 | 82.5 | 82.7 | 77.7 | 77.7 | 77.4 |
| BJNV SL4 | 88.4 | 88.5 | 88.4 | *** | 99.7 | 99.8 | 99.7 | 99.5 | 99.7 | 98.6 | 99.1 | 99.1 | 99.1 | 97.7 | 98.2 | 98.1 | 97.4 | 77.7 | 77.7 | 77.4 |
| BJNV SL3 | 88.4 | 88.5 | 88.4 | 99.8 | *** | 99.8 | 99.7 | 99.5 | 99.7 | 98.6 | 99.1 | 99 | 99 | 97.7 | 98.1 | 98.1 | 97.4 | 77.7 | 77.7 | 77.5 |
| BJNV TH3 | 88.4 | 88.5 | 88.4 | 99.9 | 99.8 | *** | 99.8 | 99.5 | 99.7 | 98.7 | 99.1 | 99.1 | 99.1 | 97.8 | 98.2 | 98.1 | 97.3 | 77.6 | 77.7 | 77.4 |
| BJNV TH4 | 88.4 | 88.5 | 88.4 | 99.9 | 99.8 | 99.9 | *** | 99.6 | 99.7 | 98.7 | 99.2 | 99.1 | 99.1 | 97.8 | 98.2 | 98.1 | 97.4 | 77.7 | 77.7 | 77.5 |
| BJNV YC4 | 88.5 | 88.5 | 88.5 | 99.8 | 99.7 | 99.8 | 99.8 | *** | 99.6 | 98.7 | 99.3 | 99.1 | 99.2 | 97.8 | 98.1 | 98.1 | 97.4 | 77.7 | 77.7 | 77.4 |
| BJNV YC3 | 88.5 | 88.5 | 88.4 | 99.9 | 99.8 | 99.9 | 99.8 | 99.8 | *** | 98.8 | 99.2 | 99.2 | 99.2 | 97.9 | 98.2 | 98.2 | 97.5 | 77.7 | 77.7 | 77.5 |
| BJNV DH3 | 88.5 | 88.7 | 88.5 | 98.6 | 98.5 | 98.5 | 98.5 | 98.5 | 98.7 | *** | 98.4 | 98.3 | 98.3 | 98 | 97.6 | 97.5 | 97.5 | 77.6 | 77.6 | 77.4 |
| BJNV H1603 | 88.3 | 88.4 | 88.3 | 99.4 | 99.3 | 99.4 | 99.4 | 99.5 | 99.5 | 98.3 | *** | 99.2 | 99.9 | 97.8 | 98 | 98 | 97 | 77.6 | 77.6 | 77.3 |
| BJNV H160 | 88.2 | 88.3 | 88.2 | 99.3 | 99.3 | 99.3 | 99.3 | 99.3 | 99.3 | 98.2 | 99.4 | *** | 99.1 | 97.7 | 97.9 | 97.9 | 97 | 77.5 | 77.5 | 77.2 |
| BJNV H801 | 88.2 | 88.3 | 88.2 | 99.3 | 99.2 | 99.3 | 99.3 | 99.4 | 99.4 | 98.2 | 99.9 | 99.3 | *** | 97.7 | 97.9 | 97.9 | 97 | 77.5 | 77.6 | 77.3 |
| BJNV H39 | 88.2 | 88.3 | 88.2 | 98.5 | 98.5 | 98.5 | 98.5 | 98.5 | 98.5 | 98.2 | 98.6 | 98.5 | 98.5 | *** | 97.2 | 97.2 | 96.9 | 77.4 | 77.4 | 77.2 |
| BJNV H56 | 87.8 | 87.8 | 87.7 | 98.5 | 98.5 | 98.6 | 98.5 | 98.5 | 98.6 | 97.5 | 98.4 | 98.5 | 98.3 | 97.8 | *** | 99.9 | 96.6 | 77.4 | 77.4 | 77.2 |
| BJNV H59 | 87.7 | 87.8 | 87.7 | 98.4 | 98.4 | 98.4 | 98.4 | 98.4 | 98.5 | 97.4 | 98.4 | 98.3 | 98.3 | 97.8 | 99.9 | *** | 96.5 | 77.4 | 77.4 | 77.1 |
| GKTV | 88.3 | 88.4 | 88.3 | 98.2 | 98.3 | 98.2 | 98.2 | 98.3 | 98.3 | 98 | 97.9 | 97.8 | 97.8 | 97.7 | 97.1 | 97 | *** | 77.6 | 77.6 | 77.4 |
| PTV | 84.8 | 84.9 | 84.8 | 84 | 84.1 | 84 | 84 | 84 | 84.1 | 84.1 | 83.9 | 83.8 | 83.8 | 83.6 | 83.3 | 83.3 | 83.9 | *** | 97.4 | 98.8 |
| NWNV1 | 84.7 | 84.8 | 84.8 | 83.8 | 83.9 | 83.8 | 83.8 | 83.8 | 83.9 | 83.9 | 83.7 | 83.7 | 83.7 | 83.4 | 83.1 | 83.1 | 83.8 | 98.3 | *** | 96.6 |
| GTV | 84.3 | 84.4 | 84.3 | 83.5 | 83.5 | 83.5 | 83.5 | 83.5 | 83.5 | 83.5 | 83.3 | 83.2 | 83.2 | 83 | 82.8 | 82.8 | 83.3 | 98.6 | 97.3 | *** |

* Abbreviations: YCNV, Yichun nariovirus; BJNV, Beiji nariovirus; GKTV, Gakugsa tick virus; PTV, Pustyn virus; NWNV1, Norway nairovirus 1; GTV, Grotenhout virus.

Table S8. Nucleotide sequence identities of S segment (upper right) of YCNV and BJNV.

|  | YCNV YC4 | YCNV YC3 | YCNV FZ3 | BJNV SL4 | BJNV SL3 | BJNV TH3 | BJNV TH4 | BJNV YC4 | BJNV YC3 | BJNV DH3 | BJNV YKS44 | BJNV H1603 | BJNV H160 | BJNV H801 | BJNV H39 | BJNV H56 | BJNV H59 | GKTV | PTV | NWNV1 | GTV |
| --- | --- | --- | --- | --- | --- | --- | --- | --- | --- | --- | --- | --- | --- | --- | --- | --- | --- | --- | --- | --- | --- |
| YCNV YC4 | *** | 98.7 | 98.7 | 83.3 | 83 | 83.2 | 83.2 | 83.2 | 83.2 | 83.2 | 83.4 | 83.6 | 83.3 | 83 | 83 | 82.7 | 82.3 | 83.5 | 76.1 | 79.4 | 78.9 |
| YCNV YC3 | *** | *** | 98.7 | 83.5 | 83.3 | 83.5 | 83.5 | 83.5 | 83.5 | 83.4 | 83.6 | 83.9 | 83.5 | 83.2 | 83.2 | 82.9 | 82.6 | 83.8 | 76 | 79.4 | 78.8 |
| YCNV FZ3 | *** | *** | *** | 83.5 | 83.3 | 83.5 | 83.5 | 83.3 | 83.3 | 83.4 | 83.5 | 83.8 | 83.5 | 83.2 | 83.2 | 82.9 | 82.4 | 83.8 | 76.2 | 79.6 | 79 |
| BJNV SL4 | *** | *** | *** | *** | 99.4 | 99.9 | 99.9 | 99.7 | 99.6 | 97.6 | 99.4 | 98.6 | 99 | 99 | 99 | 98.6 | 98.3 | 98.4 | 76.3 | 79.3 | 79.2 |
| BJNV SL3 | *** | *** | *** | *** | *** | 99.5 | 99.5 | 99.3 | 99.5 | 97.4 | 98.9 | 98.4 | 98.9 | 98.9 | 98.9 | 98.1 | 97.8 | 98.1 | 76.3 | 79.3 | 79.2 |
| BJNV TH3 | *** | *** | *** | *** | *** | *** | 100 | 99.8 | 99.7 | 97.7 | 99.5 | 98.6 | 99.1 | 99.1 | 99.1 | 98.6 | 98.4 | 98.3 | 76.3 | 79.3 | 79.2 |
| BJNV TH4 | *** | *** | *** | *** | *** | *** | *** | 99.8 | 99.7 | 97.7 | 99.5 | 98.6 | 99.1 | 99.1 | 99.1 | 98.6 | 98.4 | 98.3 | 76.3 | 79.3 | 79.2 |
| BJNV YC4 | *** | *** | *** | *** | *** | *** | *** | *** | 99.8 | 97.7 | 99.6 | 98.7 | 99 | 99 | 99 | 98.7 | 98.5 | 98.2 | 76.3 | 79.3 | 79.2 |
| BJNV YC3 | *** | *** | *** | *** | *** | *** | *** | *** | *** | 97.7 | 99.4 | 98.8 | 99.2 | 99.2 | 99.2 | 98.6 | 98.3 | 98.4 | 76.3 | 79.3 | 79.3 |
| BJNV DH3 | *** | *** | *** | *** | *** | *** | *** | *** | *** | *** | 97.3 | 97.7 | 97.3 | 97.4 | 97.4 | 96.6 | 96.2 | 96.8 | 76.1 | 79 | 79.1 |
| BJNV YKS44 | *** | *** | *** | *** | *** | *** | *** | *** | *** | *** | *** | 98.4 | 98.7 | 98.6 | 98.6 | 99.2 | 98.9 | 97.9 | 76.2 | 79.2 | 79.1 |
| BJNV H1603 | *** | *** | *** | *** | *** | *** | *** | *** | *** | *** | *** | *** | 98.4 | 98.4 | 98.4 | 97.7 | 97.4 | 98 | 76 | 79 | 79 |
| BJNV H160 | *** | *** | *** | *** | *** | *** | *** | *** | *** | *** | *** | *** | *** | 98.9 | 98.9 | 97.8 | 97.6 | 98.1 | 76.2 | 79.2 | 79.1 |
| BJNV H801 | *** | *** | *** | *** | *** | *** | *** | *** | *** | *** | *** | *** | *** | *** | 99.8 | 97.8 | 97.5 | 98.1 | 76 | 79 | 78.9 |
| BJNV H39 | *** | *** | *** | *** | *** | *** | *** | *** | *** | *** | *** | *** | *** | *** | *** | 97.8 | 97.5 | 98.2 | 76.1 | 79.1 | 79 |
| BJNV H56 | *** | *** | *** | *** | *** | *** | *** | *** | *** | *** | *** | *** | *** | *** | *** | *** | 99.3 | 97.1 | 75.6 | 78.6 | 78.5 |
| BJNV H59 | *** | *** | *** | *** | *** | *** | *** | *** | *** | *** | *** | *** | *** | *** | *** | *** | *** | 96.8 | 75.4 | 78.3 | 78.3 |
| GKTV | *** | *** | *** | *** | *** | *** | *** | *** | *** | *** | *** | *** | *** | *** | *** | *** | *** | *** | 76 | 79 | 79 |
| PTV | *** | *** | *** | *** | *** | *** | *** | *** | *** | *** | *** | *** | *** | *** | *** | *** | *** | *** | *** | 95.1 | 95.9 |
| NWNV1 | *** | *** | *** | *** | *** | *** | *** | *** | *** | *** | *** | *** | *** | *** | *** | *** | *** | *** | *** | *** | 98.6 |
| GTV | *** | *** | *** | *** | *** | *** | *** | *** | *** | *** | *** | *** | *** | *** | *** | *** | *** | *** | *** | *** | *** |

* Abbreviations: YCNV, Yichun nariovirus; BJNV, Beiji nariovirus; GKTV, Gakugsa tick virus; PTV, Pustyn virus; NWNV1, Norway nairovirus 1; GTV, Grotenhout virus.

Table S9. Nucleotide sequence identities of L segment (upper right) and amino acid sequence identities of RdRp (lower left) of MKV and MJPV*.

|  | MKV MKW73 | MKV TH3 | MKV YC4 | MKV FZ2 | MKV FZ3 | MKV ShL2 | MKV DH3 | KYV CZCT80Q | MJPV FZ3 | MJPV MDJ2 |
| --- | --- | --- | --- | --- | --- | --- | --- | --- | --- | --- |
| MKV MKW73 | *** | 93.1 | 92.5 | 92.4 | 92.4 | 92.5 | 92.9 | 83 | 76.2 | 76.1 |
| MKV TH3 | 99.1 | *** | 95.8 | 95.7 | 95.7 | 95.8 | 94.3 | 83.2 | 75.9 | 76 |
| MKV YC4 | 99.1 | 99.2 | *** | 96.2 | 96.2 | 96.3 | 94.3 | 82.7 | 76.1 | 75.9 |
| MKV FZ2 | 99.1 | 99.2 | 99.6 | *** | 100 | 98.3 | 95.4 | 82.5 | 76 | 76 |
| MKV FZ3 | 99.1 | 99.2 | 99.6 | 100 | *** | 98.3 | 95.4 | 82.5 | 76 | 76 |
| MKV ShL2 | 99 | 99.1 | 99.5 | 99.7 | 99.7 | *** | 95.7 | 82.6 | 75.9 | 75.8 |
| MKV DH3 | 99 | 98.9 | 99.1 | 99.3 | 99.3 | 99.4 | *** | 82.8 | 76.1 | 76.2 |
| KYV CZCT80Q | 95.3 | 95.3 | 95.3 | 95.3 | 95.3 | 95.2 | 95.1 | *** | 77.4 | 77.3 |
| MJPV FZ3 | 88.7 | 88.8 | 88.7 | 88.7 | 88.7 | 88.6 | 88.6 | 88.2 | *** | 95.2 |
| MJPV MDJ2 | 88.9 | 89.1 | 88.9 | 88.9 | 88.9 | 88.8 | 88.8 | 88.4 | 99.5 | *** |

* Abbreviations: MKV, Mukawa virus; KYV, Kuriyama virus; MJPV, Mudanjiang phlebovirus.

Table S10. Nucleotide sequence identities of S segment (upper right) and M segment (lower left) of MKV and MJPV*.

|  | MKV MKW73 | MKV TH3 | MKV YC4 | MKV FZ2 | MKV FZ3 | MKV ShL2 | MKV DH3 | KYV CZCT80Q | MJPV FZ3 | MJPV MDJ2 |
| --- | --- | --- | --- | --- | --- | --- | --- | --- | --- | --- |
| MKV MKW73 | *** | 93.7 | 93.5 | 93.5 | 93.5 | 93.6 | 94 | 80.1 | 70 | 69.8 |
| MKV TH3 | 90.7 | *** | 95.1 | 95 | 95.1 | 96.6 | 94.9 | 80.7 | 70.8 | 70.6 |
| MKV YC4 | 90.9 | 96.1 | *** | 95.8 | 95.9 | 94.8 | 96.8 | 80.3 | 71.1 | 70.4 |
| MKV FZ2 | 92.3 | 91.5 | 91.8 | *** | 99.9 | 94.6 | 95.1 | 80.3 | 70.6 | 70.2 |
| MKV FZ3 | 92.3 | 91.4 | 91.8 | 100 | *** | 94.7 | 95.2 | 80.3 | 70.7 | 70.3 |
| MKV ShL2 | 92.1 | 91.3 | 91.6 | 96.3 | 96.3 | *** | 95.2 | 80.5 | 70.5 | 70.4 |
| MKV DH3 | 92.2 | 91.6 | 92 | 98.3 | 98.3 | 96.8 | *** | 79.5 | 70.5 | 70.3 |
| KYV CZCT80Q | 83.5 | 84.1 | 84.4 | 84.3 | 84.3 | 83.7 | 84.1 | *** | 70.4 | 70.8 |
| MJPV FZ3 | 67.6 | 67.6 | 67.8 | 68.1 | 68.1 | 67.6 | 68 | 68.7 | *** | 96.2 |
| MJPV MDJ2 | 67.6 | 67.8 | 67.8 | 68.2 | 68.2 | 67.8 | 68.1 | 68.4 | 98.8 | *** |

* Abbreviations: MKV, Mukawa virus; KYV, Kuriyama virus; MJPV, Mudanjiang phlebovirus.

Table S11. Nucleotide sequence identities of L segment (upper right) and amino acid sequence identities of RdRp (lower left) of STPV and OTPV*.

|  | BTPV1 | NWPV1 | STPV Russia | STPV SL3 | STPV SL4 | STPV TH4 | STPV TH3 | STPV YC4 | STPV YC3 | OTPV Russia | OTPV TH4 | OTPV TH3 | OTPV SL3 | OTPV SL4 | OTPV YC3 | OTPV YC4 |
| --- | --- | --- | --- | --- | --- | --- | --- | --- | --- | --- | --- | --- | --- | --- | --- | --- |
| BTPV1 | *** | 70.6 | 70.1 | 70.2 | 70.1 | 70.1 | 70.1 | 70.3 | 70.2 | 55.9 | 55.9 | 55.7 | 55.9 | 55.8 | 55.8 | 55.9 |
| NWPV1 | 74.4 | *** | 78 | 77.9 | 77.9 | 78 | 77.9 | 78 | 77.9 | 56.7 | 56.8 | 56.8 | 56.7 | 56.7 | 56.8 | 56.8 |
| STPV Russia | 73.2 | 87 | *** | 98.1 | 98.1 | 98.1 | 98.1 | 98 | 98 | 57 | 56.8 | 56.8 | 56.8 | 56.8 | 56.9 | 56.8 |
| STPV SL3 | 73.2 | 87.3 | 99.4 | *** | 99.8 | 99.9 | 99.8 | 99.4 | 99.4 | 57 | 56.7 | 56.8 | 56.8 | 56.7 | 56.8 | 56.8 |
| STPV SL4 | 73.2 | 87.3 | 99.4 | 100 | *** | 99.9 | 99.9 | 99.4 | 99.4 | 57 | 56.8 | 56.8 | 56.8 | 56.7 | 56.8 | 56.8 |
| STPV TH4 | 73.2 | 87.3 | 99.4 | 100 | 100 | *** | 99.9 | 99.4 | 99.5 | 57 | 56.7 | 56.8 | 56.8 | 56.7 | 56.8 | 56.8 |
| STPV TH3 | 73.2 | 87.3 | 99.4 | 100 | 100 | 100 | *** | 99.4 | 99.4 | 57 | 56.7 | 56.8 | 56.8 | 56.7 | 56.8 | 56.8 |
| STPV YC4 | 73.2 | 87.4 | 99.5 | 99.9 | 99.8 | 99.9 | 99.9 | *** | 99.4 | 57 | 56.8 | 56.8 | 56.8 | 56.7 | 56.8 | 56.8 |
| STPV YC3 | 73.2 | 87.3 | 99.5 | 100 | 99.9 | 100 | 100 | 99.9 | *** | 57.1 | 56.8 | 56.8 | 56.8 | 56.8 | 56.9 | 56.8 |
| OTPV Russia | 50 | 50.2 | 50.6 | 50.9 | 50.9 | 50.9 | 50.9 | 50.8 | 50.8 | *** | 99.1 | 98.6 | 98.9 | 98.8 | 98.7 | 98.9 |
| OTPV TH4 | 50 | 50.3 | 50.7 | 50.9 | 50.9 | 50.9 | 50.9 | 50.9 | 50.9 | 99.8 | *** | 99.1 | 99.7 | 99.6 | 99.5 | 99.4 |
| OTPV TH3 | 50 | 50.2 | 50.6 | 50.9 | 50.9 | 50.9 | 50.9 | 50.8 | 50.8 | 99.8 | 99.9 | *** | 99 | 98.9 | 98.8 | 98.9 |
| OTPV SL3 | 50 | 50.3 | 50.7 | 50.9 | 50.9 | 50.9 | 50.9 | 50.9 | 50.9 | 99.8 | 99.9 | 99.9 | *** | 99.9 | 99.6 | 99.4 |
| OTPV SL4 | 50 | 50.2 | 50.6 | 50.8 | 50.9 | 50.8 | 50.8 | 50.8 | 50.8 | 99.6 | 99.7 | 99.7 | 99.8 | *** | 99.5 | 99.3 |
| OTPV YC3 | 50 | 50.3 | 50.7 | 50.9 | 50.9 | 50.9 | 50.9 | 50.9 | 50.9 | 99.6 | 99.7 | 99.7 | 99.8 | 99.6 | *** | 99.4 |
| OTPV YC4 | 50 | 50.2 | 50.6 | 50.9 | 50.9 | 50.9 | 50.9 | 50.8 | 50.8 | 99.6 | 99.7 | 99.7 | 99.8 | 99.6 | 99.9 | *** |

* Abbreviations: BTPV1, Blacklegged tick phlebovirus-1; NWPV1, Norway phlebovirus 1; STPV, Sara tick phlebovirus; OTPV, Onega tick phlebovirus.

Table S12. Nucleotide sequence identities of S segment (upper right) of STPV and OTPV*.

|  | BTPV1 | NWPV1 | STPV Russia | STPV SL3 | STPV SL4 | STPV TH4 | STPV TH3 | STPV YC4 | STPV YC3 | OTPV Russia | OTPV TH4 | OTPV TH3 | OTPV SL3 | OTPV SL4 | OTPV YC3 | OTPV YC4 |
| --- | --- | --- | --- | --- | --- | --- | --- | --- | --- | --- | --- | --- | --- | --- | --- | --- |
| BTPV1 | *** | 53.3 | 54.8 | 55.1 | 55 | 55 | 55 | 55 | 55 | 38.9 | 39.1 | 39 | 39 | 39 | 38.9 | 39.2 |
| NWPV1 | *** | *** | 83.8 | 83.6 | 83.5 | 83.5 | 83.5 | 83.5 | 83.5 | 53.6 | 53.8 | 53.7 | 53.6 | 53.8 | 53.6 | 53.4 |
| STPV Russia | *** | *** | *** | 99.2 | 99.1 | 99.1 | 99 | 99 | 99 | 56.4 | 56.8 | 56.9 | 56.7 | 56.8 | 57 | 56.8 |
| STPV SL3 | *** | *** | *** | *** | 99.8 | 99.8 | 99.9 | 99.8 | 99.8 | 56.8 | 57.2 | 57.3 | 57.1 | 57.2 | 57.3 | 57.1 |
| STPV SL4 | *** | *** | *** | *** | *** | 100 | 99.9 | 99.6 | 99.6 | 56.7 | 57.1 | 57.2 | 57 | 57.1 | 57.3 | 57.1 |
| STPV TH4 | *** | *** | *** | *** | *** | *** | 99.9 | 99.6 | 99.6 | 56.7 | 57.1 | 57.2 | 57 | 57.1 | 57.3 | 57.1 |
| STPV TH3 | *** | *** | *** | *** | *** | *** | *** | 99.6 | 99.6 | 56.7 | 57.1 | 57.2 | 57 | 57.1 | 57.3 | 57.1 |
| STPV YC4 | *** | *** | *** | *** | *** | *** | *** | *** | 100 | 56.8 | 57.2 | 57.3 | 57.1 | 57.2 | 57.3 | 57.2 |
| STPV YC3 | *** | *** | *** | *** | *** | *** | *** | *** | *** | 56.8 | 57.2 | 57.3 | 57.1 | 57.2 | 57.3 | 57.2 |
| OTPV Russia | *** | *** | *** | *** | *** | *** | *** | *** | *** | *** | 98.8 | 98.8 | 98.8 | 98.4 | 98.6 | 98.5 |
| OTPV TH4 | *** | *** | *** | *** | *** | *** | *** | *** | *** | *** | *** | 99.5 | 99.9 | 99.3 | 99 | 98.8 |
| OTPV TH3 | *** | *** | *** | *** | *** | *** | *** | *** | *** | *** | *** | *** | 99.5 | 99.4 | 99.5 | 99.3 |
| OTPV SL3 | *** | *** | *** | *** | *** | *** | *** | *** | *** | *** | *** | *** | *** | 99.2 | 99 | 98.8 |
| OTPV SL4 | *** | *** | *** | *** | *** | *** | *** | *** | *** | *** | *** | *** | *** | *** | 99 | 98.8 |
| OTPV YC3 | *** | *** | *** | *** | *** | *** | *** | *** | *** | *** | *** | *** | *** | *** | *** | 99.5 |
| OTPV YC4 | *** | *** | *** | *** | *** | *** | *** | *** | *** | *** | *** | *** | *** | *** | *** | *** |

* Abbreviations: BTPV1, Blacklegged tick phlebovirus-1; NWPV1, Norway phlebovirus 1; STPV, Sara tick phlebovirus; OTPV, Onega tick phlebovirus.

Table S13. Nucleotide sequence identities of complete genome (upper right) and amino acid sequence identities of RdRp (lower left) of THRV1, THRV2, THRV3*.

|  | THRV2 TH3 | THRV2 TH4 | THRV3 TH3 | THRV1 TH1 | THRV1 TH2 | THRV1 SL1 | THRV1 SL2 | THRV1 DH1 | THRV1 YC2 | THRV1 ShL3 | THRV1 JA | WVMV1 | MLV | HPTV3 | BLTV2 | TCTV3 | WHTV1 |
| --- | --- | --- | --- | --- | --- | --- | --- | --- | --- | --- | --- | --- | --- | --- | --- | --- | --- |
| THRV2 TH3 | *** | 98.9 | 45.9 | 48.7 | 48.6 | 48.7 | 48.6 | 49.4 | 49.3 | 49.3 | 49.3 | 69 | 35.8 | 45.5 | 45.6 | 44.4 | 44.9 |
| THRV2 TH4 | 99.6 | *** | 46.1 | 48.7 | 48.6 | 48.7 | 48.6 | 49.3 | 49.3 | 49.3 | 49.2 | 69.1 | 35.9 | 45.6 | 45.6 | 44.3 | 44.9 |
| THRV3 TH3 | 47.7 | 47.7 | *** | 44.9 | 44.9 | 44.9 | 44.9 | 44.9 | 44.9 | 44.9 | 44.9 | 45.9 | 32.1 | 43 | 42.4 | 45.9 | 43.8 |
| THRV1 TH1 | 50.1 | 50.1 | 44.4 | *** | 99.3 | 100 | 99.3 | 81.4 | 81.4 | 81.4 | 81.4 | 49 | 30.6 | 45.3 | 49.3 | 50.1 | 50.6 |
| THRV1 TH2 | 50.1 | 50.1 | 44.4 | 100 | *** | 99.3 | 100 | 81.4 | 81.4 | 81.4 | 81.4 | 49 | 30.6 | 45.4 | 49.4 | 50.1 | 50.5 |
| THRV1 SL1 | 50.1 | 50.1 | 44.4 | 100 | 100 | *** | 99.3 | 81.4 | 81.4 | 81.4 | 81.4 | 49 | 30.6 | 45.3 | 49.3 | 50.1 | 50.6 |
| THRV1 SL2 | 50.1 | 50.1 | 44.4 | 100 | 100 | 100 | *** | 81.4 | 81.4 | 81.4 | 81.4 | 49 | 30.6 | 45.4 | 49.4 | 50.1 | 50.5 |
| THRV1 DH1 | 50.6 | 50.6 | 44.4 | 93.7 | 93.7 | 93.7 | 93.7 | *** | 99.7 | 99.4 | 99.5 | 48.8 | 30.5 | 45.3 | 49 | 50 | 50.5 |
| THRV1 YC2 | 50.6 | 50.6 | 44.4 | 93.7 | 93.7 | 93.7 | 93.7 | 100 | *** | 99.1 | 99.2 | 48.8 | 30.6 | 45.3 | 49 | 50 | 50.6 |
| THRV1 ShL3 | 50.6 | 50.6 | 44.4 | 93.7 | 93.7 | 93.7 | 93.7 | 99.9 | 99.9 | *** | 99.3 | 48.8 | 30.5 | 45.3 | 49.1 | 49.9 | 50.6 |
| THRV1 JA | 50.6 | 50.6 | 44.4 | 93.6 | 93.6 | 93.6 | 93.6 | 99.7 | 99.7 | 99.7 | *** | 48.8 | 30.4 | 45.3 | 49 | 50 | 50.5 |
| NWMV1 | 79.8 | 79.7 | 48.1 | 51.2 | 51.2 | 51.2 | 51.2 | 51 | 51 | 51 | 51 | *** | 35.5 | 46 | 45.8 | 44.9 | 45.2 |
| MLV | 50.7 | 50.7 | 44.4 | 70.7 | 70.7 | 70.7 | 70.7 | 70.3 | 70.3 | 70.3 | 70.2 | 50.9 | *** | 30.1 | 29.9 | 29.1 | 29.7 |
| HPTV3 | 46.8 | 46.9 | 43.8 | 43.7 | 43.7 | 43.7 | 43.7 | 43.7 | 43.7 | 43.7 | 43.7 | 47.3 | 44.7 | *** | 44 | 43.6 | 44.2 |
| BLTV2 | 45.9 | 45.9 | 41.4 | 49.9 | 49.9 | 49.9 | 49.9 | 49.9 | 49.9 | 49.9 | 49.9 | 46.1 | 50.7 | 41.7 | *** | 46.2 | 46.5 |
| TCTV3 | 45.7 | 45.7 | 43 | 52.3 | 52.3 | 52.3 | 52.3 | 51.8 | 51.8 | 51.8 | 51.8 | 46.8 | 51.5 | 43 | 49 | *** | 58.3 |
| WHTV1 | 45.2 | 45.3 | 41.2 | 49.5 | 49.5 | 49.5 | 49.5 | 49.5 | 49.5 | 49.5 | 49.5 | 45.4 | 50.4 | 41.9 | 47.1 | 64.3 | *** |

* Abbreviations: THRV1, Tahe rhabdovirus 1; THRV2, Tahe rhabdovirus 2; THRV3, Tahe rhabdovirus 3; OTPV, Onega tick phlebovirus; NWMV1: Norway mononegavirus 1; MLV: Manly virus; HPTV3: Huangpi tick virus 3; BLTV2: Bole tick virus 2; Tacheng tick virus 3: TCTV3; WHTV1: Wuhan tick virus 1.

Table S14. Nucleotide sequence identities of complete genome (upper right) and amino acid sequence identities of RdRp (lower left) of NUMV*.

|  | DTMV | SFV | LMV | H109 | H43 | H141 | T43 | H159 | H145 | H160 | NE-MDJ2 | NE-FZ3 | NE-FZ4 | NE-SL3 | NE-SL4 | NE-TH4 | NE-YC4 | NE-YC3 | NE-TH3 | NE-DH3 |
| --- | --- | --- | --- | --- | --- | --- | --- | --- | --- | --- | --- | --- | --- | --- | --- | --- | --- | --- | --- | --- |
| DTMV | *** | 99.9 | 64.8 | 64.9 | 65 | 64.8 | 64.9 | 64.6 | 64.5 | 64.7 | 64.9 | 64.9 | 64.9 | 65 | 65 | 65 | 64.9 | 64.9 | 64.9 | 64.9 |
| SFV | 99.9 | *** | 64.8 | 64.9 | 65.1 | 64.9 | 64.9 | 64.7 | 64.6 | 64.8 | 65 | 64.9 | 64.9 | 65 | 65 | 65 | 65 | 64.9 | 64.9 | 64.9 |
| LMV | 76.5 | 76.4 | *** | 96.8 | 96.6 | 97.1 | 97.5 | 96.7 | 96.7 | 96.6 | 96.6 | 96.6 | 96.6 | 98.2 | 98.3 | 96.9 | 97 | 96.7 | 96.8 | 96.8 |
| H109 | 76.6 | 76.6 | 99.4 | *** | 98.5 | 97.6 | 98.1 | 97.5 | 97.4 | 97.4 | 98.8 | 98.9 | 98.9 | 97.9 | 97.8 | 98.9 | 98.7 | 99 | 99.1 | 98.2 |
| H43 | 76.6 | 76.6 | 99.4 | 99.9 | *** | 97.6 | 97.4 | 97.4 | 97.4 | 97.3 | 98.4 | 98.4 | 98.4 | 97.6 | 97.5 | 98.5 | 98.5 | 98.5 | 98.6 | 98.3 |
| H141 | 75.9 | 75.9 | 99 | 98.7 | 98.7 | *** | 99.1 | 97.5 | 97.6 | 97.6 | 97.5 | 97.6 | 97.6 | 98.5 | 98.3 | 97.8 | 97.8 | 97.7 | 97.8 | 97.5 |
| T43 | 75.9 | 75.9 | 99 | 98.7 | 98.6 | 100 | *** | 97.6 | 97.3 | 97.3 | 97.4 | 97.5 | 97.4 | 98.9 | 98.9 | 97.7 | 97.6 | 97.6 | 97.6 | 97.3 |
| H159 | 76.5 | 76.4 | 99.2 | 99.3 | 99.4 | 98.5 | 98.5 | *** | 99.2 | 99 | 97.3 | 97.3 | 97.3 | 97.7 | 97.7 | 97.5 | 97.8 | 97.5 | 97.5 | 97.3 |
| H145 | 76.4 | 76.3 | 99.3 | 99.3 | 99.3 | 98.6 | 98.6 | 99.8 | *** | 99.1 | 97.2 | 97.2 | 97.2 | 97.7 | 97.6 | 97.5 | 97.8 | 97.4 | 97.4 | 97.2 |
| H160 | 76.4 | 76.3 | 99.4 | 99.4 | 99.3 | 98.7 | 98.6 | 99.8 | 99.8 | *** | 97.2 | 97.2 | 97.2 | 97.7 | 97.5 | 97.5 | 97.8 | 97.5 | 97.4 | 97.3 |
| NE-MDJ2 | 76.6 | 76.6 | 99.3 | 99.8 | 99.8 | 98.6 | 98.5 | 99.2 | 99.1 | 99.2 | *** | 99.5 | 99.5 | 97.5 | 97.4 | 99.5 | 99.3 | 99.6 | 99.7 | 98.6 |
| NE-FZ3 | 76.6 | 76.6 | 99.3 | 99.8 | 99.8 | 98.6 | 98.5 | 99.2 | 99.1 | 99.2 | 99.7 | *** | 100 | 97.6 | 97.5 | 99.5 | 99.3 | 99.7 | 99.7 | 98.7 |
| NE-FZ4 | 76.6 | 76.5 | 99.2 | 99.8 | 99.8 | 98.5 | 98.5 | 99.1 | 99.1 | 99.2 | 99.7 | 99.9 | *** | 97.6 | 97.4 | 99.5 | 99.3 | 99.7 | 99.7 | 98.7 |
| NE-SL3 | 76.5 | 76.4 | 99.7 | 99.4 | 99.4 | 99.2 | 99.1 | 99.2 | 99.3 | 99.3 | 99.3 | 99.3 | 99.2 | *** | 99.7 | 97.9 | 97.9 | 97.7 | 97.8 | 97.7 |
| NE-SL4 | 76.5 | 76.4 | 99.8 | 99.4 | 99.4 | 99.3 | 99.2 | 99.3 | 99.4 | 99.4 | 99.3 | 99.3 | 99.3 | 99.9 | *** | 97.8 | 97.8 | 97.6 | 97.7 | 97.6 |
| NE-TH4 | 76.6 | 76.6 | 99.4 | 99.9 | 99.9 | 98.8 | 98.7 | 99.3 | 99.3 | 99.4 | 99.8 | 99.8 | 99.8 | 99.4 | 99.5 | *** | 99.4 | 99.6 | 99.7 | 98.8 |
| NE-YC4 | 76.6 | 76.6 | 99.3 | 99.8 | 99.8 | 98.6 | 98.6 | 99.5 | 99.4 | 99.5 | 99.7 | 99.7 | 99.6 | 99.3 | 99.4 | 99.8 | *** | 99.4 | 99.5 | 98.9 |
| NE-YC3 | 76.6 | 76.6 | 99.4 | 99.9 | 99.8 | 98.8 | 98.7 | 99.4 | 99.3 | 99.4 | 99.7 | 99.8 | 99.8 | 99.4 | 99.5 | 99.8 | 99.7 | *** | 99.9 | 98.8 |
| NE-TH3 | 76.6 | 76.6 | 99.4 | 99.9 | 99.9 | 98.7 | 98.7 | 99.3 | 99.3 | 99.4 | 99.8 | 99.9 | 99.8 | 99.4 | 99.4 | 99.9 | 99.7 | 100 | *** | 98.9 |
| NE-DH3 | 76.6 | 76.5 | 99.5 | 99.8 | 99.7 | 98.8 | 98.8 | 99.4 | 99.4 | 99.5 | 99.6 | 99.6 | 99.6 | 99.5 | 99.6 | 99.7 | 99.7 | 99.8 | 99.8 | *** |

* Abbreviations: NUMV, Nuomin virus; DTMV, Deer tick mononegavirales; SFV, Suffolk virus; LMV, Lesnoe mivirus. NUMV was omitted from the table and replaced by the strain name.

Table S15. Nucleotide sequence identities of RdRp genome (upper right) and amino acid sequence identities of RdRp (lower left) of JPLV1*.

|  | JPLV1 YC3 | JPLV1 YC4 | JPLV1 SL3 | JPLV1 DH3 | JPLV1 MDJ2 | JPLV1 FZ4 | JPLV1 FZ3 | JPLV1 QG-4 | JPLV1 QG-1 | JPLV1 QG-2 | NPLV1 NOR/S5/Kilen/2014 | NPLV1 NOR/H3/Kilen/2014 |
| --- | --- | --- | --- | --- | --- | --- | --- | --- | --- | --- | --- | --- |
| JPLV1 YC3 | *** | 99.5 | 99.3 | 99.7 | 99.5 | 99.4 | 99.5 | 99.7 | 99.5 | 99.5 | 91.9 | 91.6 |
| JPLV1 YC4 | 100 | *** | 99.8 | 99.7 | 99.6 | 99.7 | 99.5 | 99.5 | 99.4 | 99.4 | 91.9 | 91.5 |
| JPLV1 SL3 | 99.8 | 99.8 | *** | 99.5 | 99.4 | 99.9 | 99.6 | 99.3 | 99.3 | 99.2 | 91.9 | 91.5 |
| JPLV1 DH3 | 99.8 | 99.8 | 99.5 | *** | 99.6 | 99.4 | 99.5 | 99.5 | 99.4 | 99.4 | 91.8 | 91.3 |
| JPLV1 MDJ2 | 100 | 100 | 99.8 | 99.8 | *** | 99.3 | 99.5 | 99.5 | 99.3 | 99.3 | 91.9 | 91.5 |
| JPLV1 FZ4 | 99.8 | 99.8 | 100 | 99.5 | 99.8 | *** | 99.7 | 99.4 | 99.4 | 99.2 | 91.9 | 91.6 |
| JPLV1 FZ3 | 99.8 | 99.8 | 100 | 99.5 | 99.8 | 100 | *** | 99.5 | 99.4 | 99.4 | 91.9 | 91.6 |
| JPLV1 QG-4 | 100 | 100 | 99.8 | 99.8 | 100 | 99.8 | 99.8 | *** | 99.7 | 99.8 | 91.9 | 91.6 |
| JPLV1 QG-1 | 100 | 100 | 99.8 | 99.8 | 100 | 99.8 | 99.8 | 100 | *** | 99.8 | 91.9 | 91.8 |
| JPLV1 QG-2 | 100 | 100 | 99.8 | 99.8 | 100 | 99.8 | 99.8 | 100 | 100 | *** | 91.9 | 91.8 |
| NPLV1 NOR/S5/Kilen/2014 | 94.3 | 94.3 | 94.3 | 94.1 | 94.3 | 94.3 | 94.3 | 94.3 | 94.3 | 94.3 | *** | 98 |
| NPLV1 NOR/H3/Kilen/2014 | 94.1 | 94.1 | 94.1 | 93.9 | 94.1 | 94.1 | 94.1 | 94.1 | 94.1 | 94.1 | 98.6 | *** |

* Abbreviations: JPLV1, Jilin partiti-like virus 1; NPLV1, Norway partiti-like virus 1.

Table S16. Nucleotide sequence identities of complete cds (upper right) and amino acid sequence identities of RdRp (lower left) of FLTV*.

|  | FTLV YC3 | FTLV FZ4 | FTLV FZ3 | FTLV DH3 | UMV | STTLV |
| --- | --- | --- | --- | --- | --- | --- |
| FTLV YC3 | *** | 92.9 | 92.7 | 89.8 | 73.6 | 37 |
| FTLV FZ4 | 98.8 | *** | 93.1 | 92.9 | 73.6 | 37.2 |
| FTLV FZ3 | 94.9 | 95.8 | *** | 92.9 | 73.5 | 36.9 |
| FTLV DH3 | 95.6 | 96.5 | 97.9 | *** | 73.4 | 36.9 |
| UMV | 81.1 | 81.3 | 80.6 | 80.6 | *** | 37.8 |
| STTLV | 30.3 | 30.5 | 30 | 30 | 33.3 | *** |

* Abbreviations: FTLV, Fangzheng tombus-like virus; UMV, Upmeje virus; STTLV, Soybean thrips tombus-like virus.

Table S17. Nucleotide sequence identities of complete cds (upper right) and amino acid sequence identities of RdRp (lower left) of ISAV1 and XTAV1*.

|  | ISAV1 ISE6 | NLLV3 | ISAV1 SL4 | ISAV1 SL3 | ISAV1 TH4 | ISAV1 YC4 | ISAV1 YC3 | ISAV1 DH3 | ADTV1 | XTAV1 14-YG | XTAV1 16-T2 | XTAV1 ShL1 | XTAV1 ShL2 | XTAV1 ShL3 | XTAV1 DH2 |
| --- | --- | --- | --- | --- | --- | --- | --- | --- | --- | --- | --- | --- | --- | --- | --- |
| ISAV1 ISE6 | *** | 82.1 | 81.8 | 81.8 | 82.1 | 81.5 | 81.6 | 81.6 | 62.4 | 61.6 | 61.7 | 61.9 | 61.6 | 61.8 | 62.1 |
| NLLV3 | 88.5 | *** | 85.5 | 85.5 | 85.7 | 85.4 | 85.7 | 85.7 | 61.9 | 62.3 | 62.3 | 62.7 | 62.4 | 62.5 | 62.5 |
| ISAV1 SL4 | 87.7 | 93.4 | *** | 100 | 98.6 | 98.8 | 99.1 | 99.2 | 61.7 | 62.1 | 62 | 62.4 | 61.9 | 62.3 | 62.3 |
| ISAV1 SL3 | 87.7 | 93.4 | 100 | *** | 98.6 | 98.8 | 99.1 | 99.2 | 61.7 | 62.1 | 62 | 62.4 | 61.9 | 62.3 | 62.3 |
| ISAV1 TH4 | 87.7 | 93.4 | 100 | 100 | *** | 98.1 | 98.8 | 98.6 | 62.3 | 62.3 | 62.4 | 62.8 | 62.3 | 62.8 | 62.6 |
| ISAV1 YC4 | 87.7 | 93.4 | 100 | 100 | 100 | *** | 99 | 98.7 | 62 | 62 | 62.1 | 62.6 | 62.1 | 62.4 | 62.4 |
| ISAV1 YC3 | 87.7 | 93.4 | 100 | 100 | 100 | 100 | *** | 99 | 61.9 | 62 | 62.1 | 62.7 | 62.2 | 62.5 | 62.4 |
| ISAV1 DH3 | 87.3 | 93 | 99.6 | 99.6 | 99.6 | 99.6 | 99.6 | *** | 62 | 62.1 | 62 | 62.4 | 61.9 | 62.3 | 62.2 |
| ADTV1 | 70.5 | 70.5 | 72.1 | 72.1 | 72.1 | 72.1 | 72.1 | 71.7 | *** | 66.7 | 66.8 | 67.1 | 67.1 | 66.7 | 67 |
| XTAV1 14-YG | 67.9 | 70 | 69.1 | 69.1 | 69.1 | 69.1 | 69.1 | 68.7 | 78.2 | *** | 99.3 | 96.1 | 96.2 | 95.9 | 96.1 |
| XTAV1 16-T2 | 67.9 | 70 | 69.1 | 69.1 | 69.1 | 69.1 | 69.1 | 68.7 | 78.2 | 100 | *** | 96.2 | 96.2 | 96 | 96 |
| XTAV1 ShL1 | 67.9 | 70 | 69.1 | 69.1 | 69.1 | 69.1 | 69.1 | 68.7 | 78.2 | 99.2 | 99.2 | *** | 97.5 | 97.1 | 98.3 |
| XTAV1 ShL2 | 67.9 | 70 | 69.1 | 69.1 | 69.1 | 69.1 | 69.1 | 68.7 | 78.6 | 98.8 | 98.8 | 99.6 | *** | 98.3 | 97.8 |
| XTAV1 ShL3 | 67.9 | 70 | 69.1 | 69.1 | 69.1 | 69.1 | 69.1 | 68.7 | 77.4 | 97.9 | 97.9 | 98.8 | 98.4 | *** | 97.4 |
| XTAV1 DH2 | 67.9 | 70 | 69.1 | 69.1 | 69.1 | 69.1 | 69.1 | 68.7 | 78.6 | 98.8 | 98.8 | 99.6 | 100 | 98.4 | *** |

* Abbreviations: ISAV1, Ixodes scapularis associated virus 1; XTAV1, Xinjiang tick associated virus 1; NLLV3, Norway luteo-like virus 3; ADTV, American dog tick associated virus-1.

Table S18. Nucleotide sequence identities of complete cds (upper right) and amino acid sequence identities of RdRp (lower left) of JLLV2*.

|  | JLLV2 SL4 | JLLV2 TH3 | JLLV2 YC4 | JLLV2 YC3 | JLLV2 FZ4 | JLLV2 DH3 | JLLV2 QG-2 | JLLV2 YQG-3 | NLLV2 H3 | NLLV2 A2 |
| --- | --- | --- | --- | --- | --- | --- | --- | --- | --- | --- |
| JLLV2 SL4 | *** | 98.3 | 98.7 | 98.6 | 98.6 | 99 | 98.7 | 98.5 | 87.7 | 87.7 |
| JLLV2 TH3 | 99.5 | *** | 98.3 | 98.2 | 98.1 | 98.6 | 98.4 | 98.3 | 87.9 | 87.9 |
| JLLV2 YC4 | 99.8 | 99.8 | *** | 100 | 98.9 | 99.3 | 98.6 | 98.5 | 87.9 | 87.9 |
| JLLV2 YC3 | 99.8 | 99.8 | 100 | *** | 98.9 | 99.2 | 98.6 | 98.4 | 88 | 88 |
| JLLV2 FZ4 | 99.5 | 99.5 | 99.8 | 99.8 | *** | 99.2 | 98.7 | 98.4 | 87.9 | 87.9 |
| JLLV2 DH3 | 99.9 | 99.6 | 99.9 | 99.9 | 99.6 | *** | 99 | 98.8 | 88.2 | 88.2 |
| JLLV2 QG-2 | 99.6 | 99.6 | 99.9 | 99.9 | 99.6 | 99.8 | *** | 99.8 | 87.9 | 87.9 |
| JLLV2 YQG-3 | 99.5 | 99.8 | 99.8 | 99.8 | 99.5 | 99.6 | 99.9 | *** | 87.9 | 87.9 |
| NLLV2 H3 | 91.7 | 91.5 | 91.5 | 91.5 | 91.3 | 91.7 | 91.4 | 91.5 | *** | 100 |
| NLLV2 A2 | 91.7 | 91.5 | 91.5 | 91.5 | 91.3 | 91.7 | 91.4 | 91.5 | 100 | *** |

* Abbreviations: JLLV2, Jilin luteo-like virus 2; NLLV2, Norway luteo-like virus 2.

Table S19. Nucleotide sequence identities of complete genome (upper right) of ISAV3*.

|  | ISAV3 TH4 | ISAV3 SL4 | ISAV3 YC4 | ISAV3 YC3 | ISAV3 ISE6 | ISAV4 RTS-11 | XTAV2 15-CYFC39 | XTAV2 381unRc | ADTV2 RTS-100 | ADTV2 RTS-1100 |
| --- | --- | --- | --- | --- | --- | --- | --- | --- | --- | --- |
| ISAV3 SL3 | 94.5 | 99.8 | 94.8 | 93.7 | 87.5 | 88 | 66.4 | 65.3 | 67.9 | 67.9 |
| ISAV3 TH4 | *** | 94.6 | 99.3 | 98.9 | 87.1 | 87.3 | 66.2 | 65.3 | 67.5 | 67.5 |
| ISAV3 SL4 | *** | *** | 95 | 93.9 | 87.6 | 88.2 | 66.4 | 65.5 | 68.1 | 68.1 |
| ISAV3 YC4 | *** | *** | *** | 98.9 | 87.5 | 87.5 | 66.6 | 65.7 | 67.7 | 67.7 |
| ISAV3 YC3 | *** | *** | *** | *** | 86.7 | 86.7 | 66.4 | 66.1 | 67.9 | 67.9 |
| ISAV3 ISE6 | *** | *** | *** | *** | *** | 98.3 | 66.4 | 67.3 | 69.4 | 69.4 |
| ISAV4 RTS-11 | *** | *** | *** | *** | *** | *** | 67 | 67.3 | 69.7 | 69.7 |
| XTAV2 15-CYFC39 | *** | *** | *** | *** | *** | *** | *** | 81.9 | 65.5 | 65.5 |
| XTAV2 381unRc | *** | *** | *** | *** | *** | *** | *** | *** | 66.6 | 66.6 |
| ADTV2 RTS-100 | *** | *** | *** | *** | *** | *** | *** | *** | *** | 100 |

* Abbreviations: ISAV3, Ixodes scapularis associated virus 3; ISAV4, Ixodes scapularis associated virus 4; XTAV2, Xinjiang tick associated virus 2; ADTV, ISAV3, American dog tick associated virus 2.
