## Supplementary figures and images for "Extensive diversity of RNA viruses in ticks revealed by metagenomics in northeastern China"

### Supplementary Fig. S1.pdf

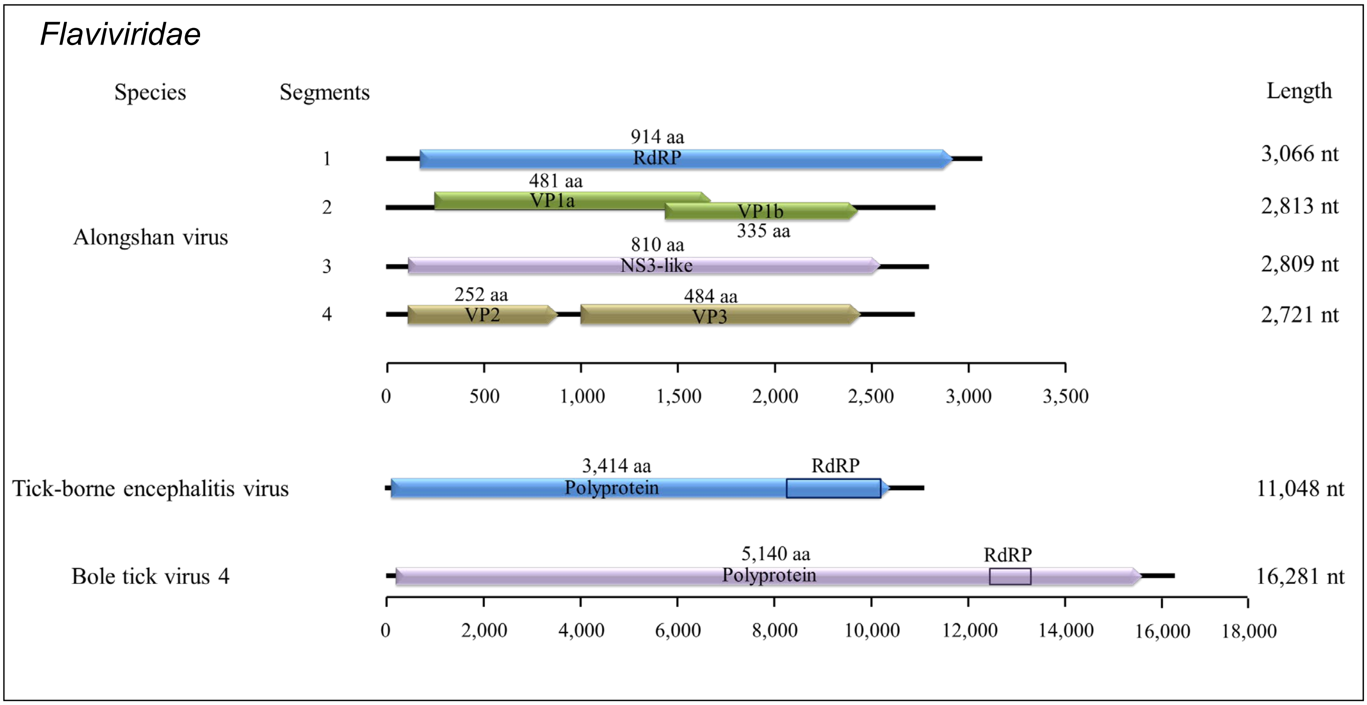

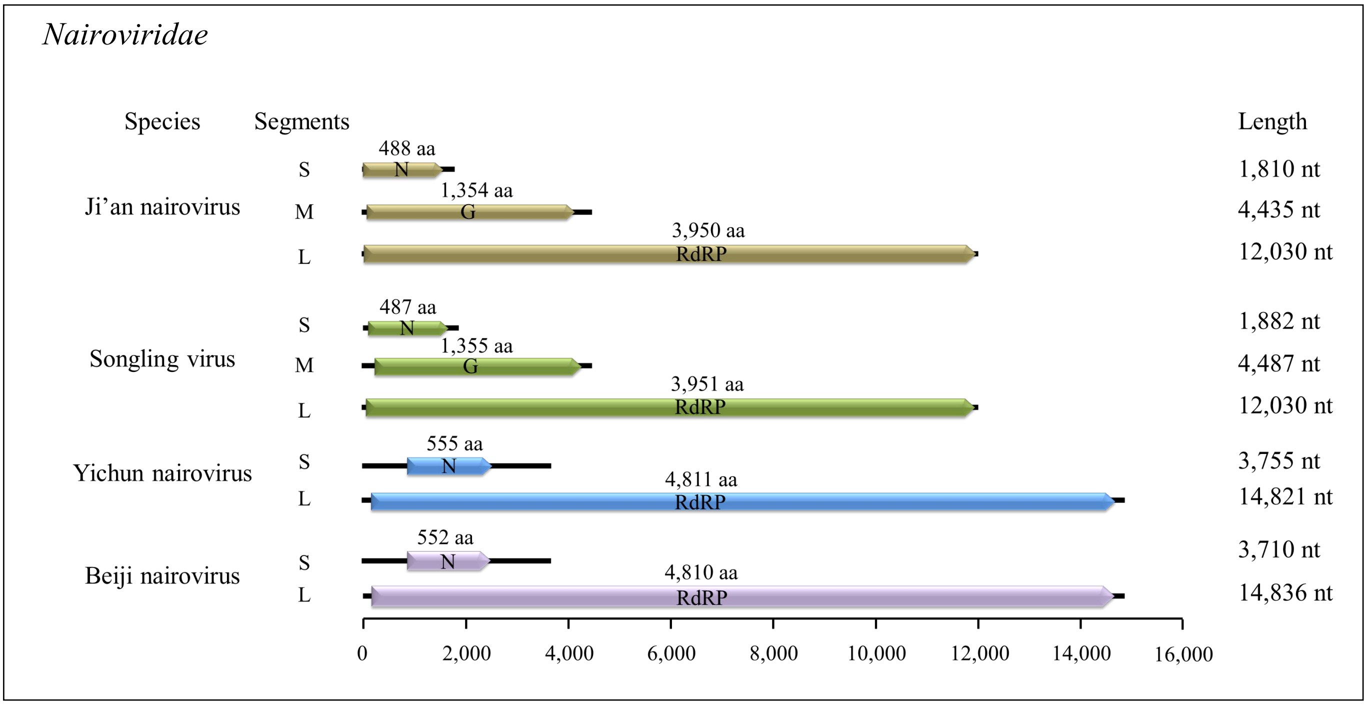

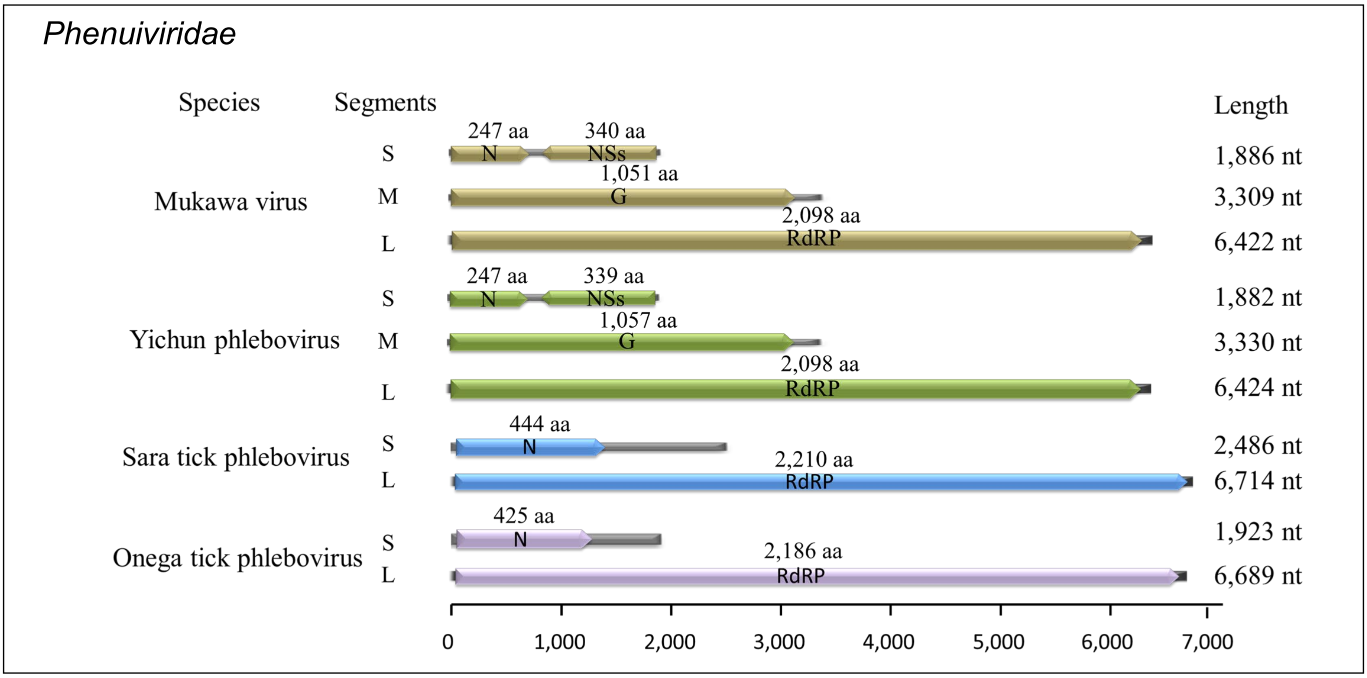

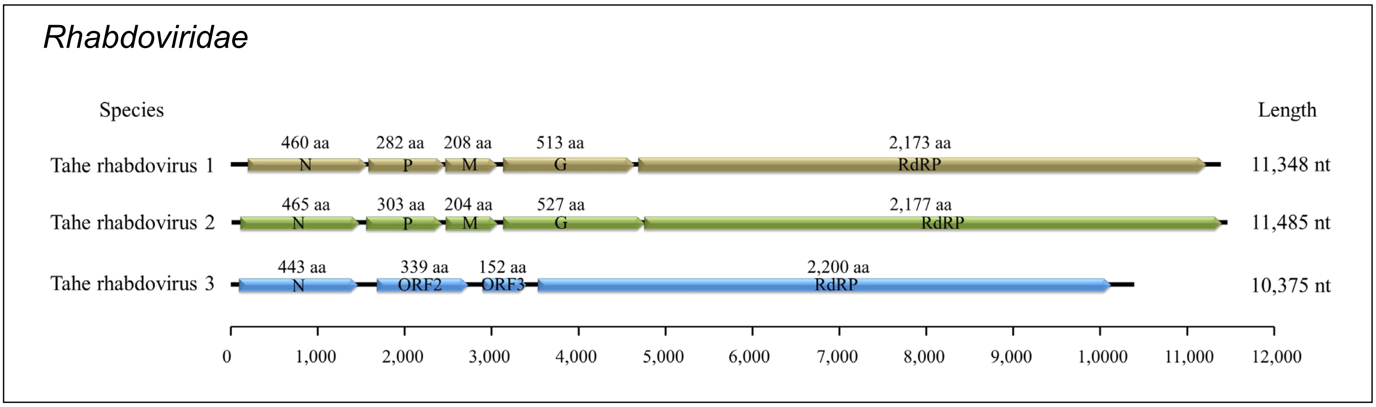

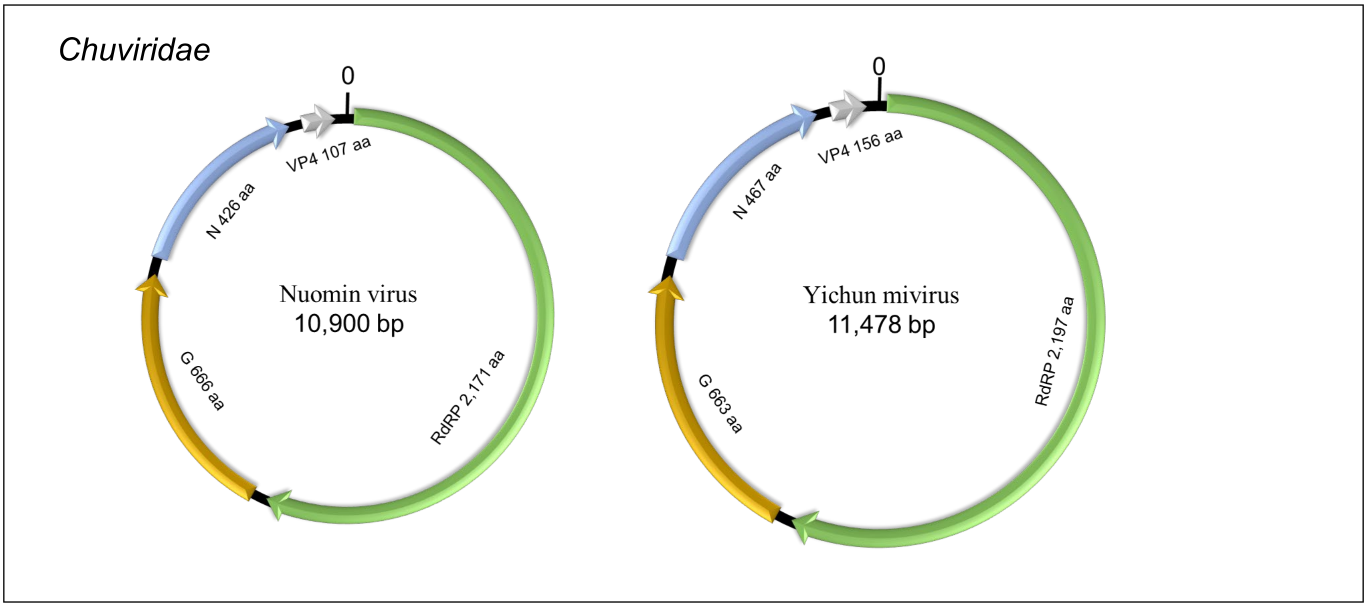

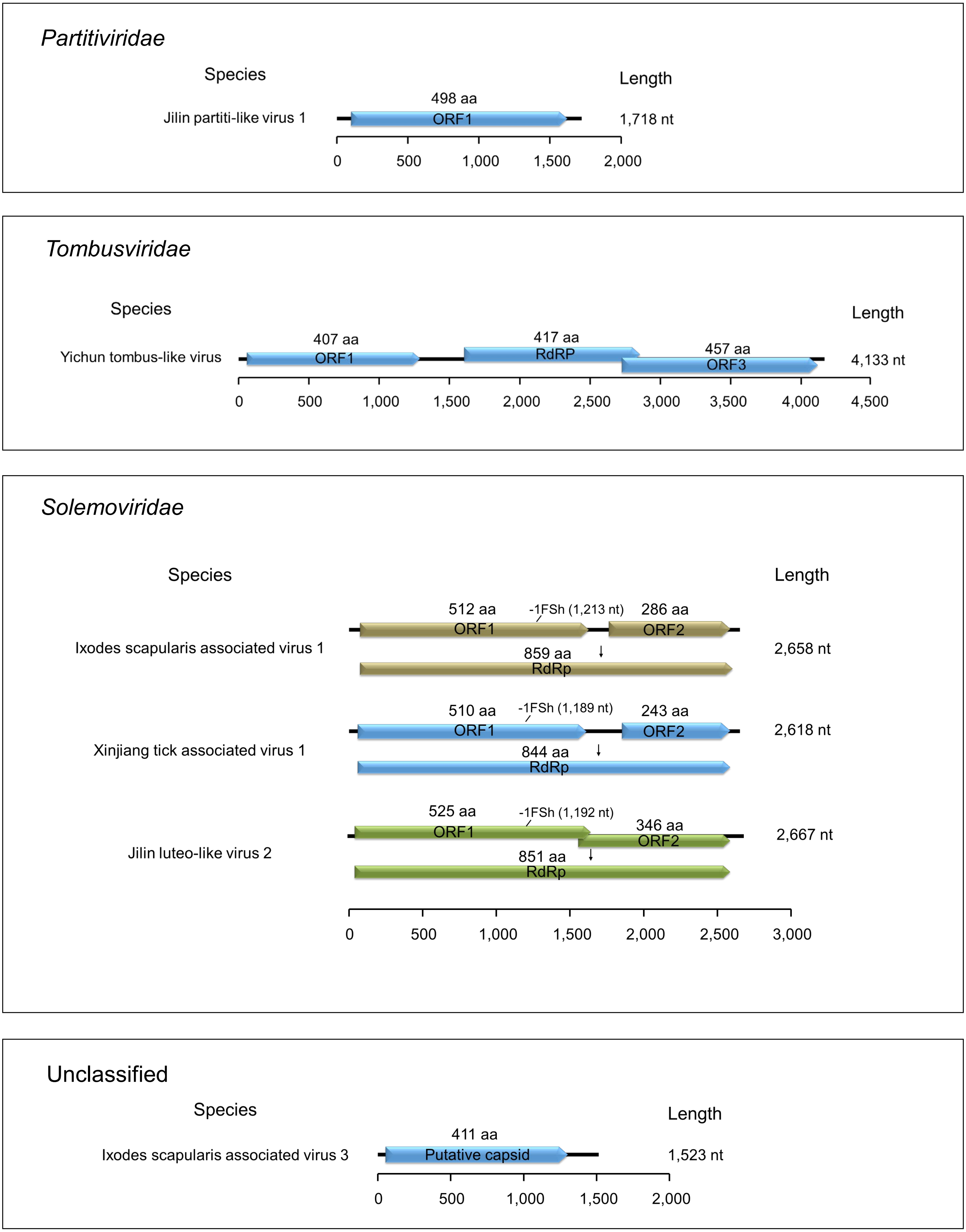
